# SELECTIVE ACTIVATION OF GIRK POTASSIUM CHANNELS REDUCES BEHAVIORAL AND BRAIN RESPONSES TO ETHANOL IN MICE

**DOI:** 10.64898/2026.01.30.702835

**Authors:** Jaume Taura, Arianna Marrazzo, Se In Son, Ganesha Rai, Max Kreifeldt, Candice Contet, Paul A. Slesinger

## Abstract

Alcohol use disorder (AUD) is a chronic relapsing condition with limited pharmacological treatments. Ethanol modulates neuronal excitability in part through activation of G-protein-gated inwardly rectifying potassium (GIRK/Kir3) channels, which dampen neuronal activity in reward- and stress-related circuits implicated in AUD pathophysiology. In this study, we investigated the therapeutic potential of targeting activation of GIRK channels in mouse models of ethanol intoxication. GiGA1 (G protein-independent GIRK activator type 1) is a selective activator of GIRK1/GIRK2 channels and has good brain bioavailability. Systemic GiGA1 administration prevented acquisition of ethanol-induced conditioned place preference (CPP) in both male and female mice. GiGA1 also significantly reduced voluntary ethanol intake and decreased blood alcohol concentrations, when administered to mice after they developed high preference and consumption of ethanol. Similarly, Baclofen, a GABA_B_ receptor agonist that leads to activation of GIRK channels also decreased ethanol consumption. However, systemic Baclofen did not prevent acquisition of ethanol-dependent CPP, suggesting a broader efficacy of direct GIRK1/GIRK2 activation by GiGA1. Whole-brain c-Fos mapping as a proxy for neuronal activity revealed that GiGA1 blunted ethanol-induced neuronal activation in several AUD-relevant brain regions, including the central amygdala, paraventricular thalamus, paraventricular hypothalamus, and Edinger–Westphal nucleus. These findings demonstrate that pharmacological activation of GIRK channels modulates key neural circuits involved in ethanol reward and intake, supporting GiGA1 as a promising lead compound for targeted AUD therapy.

## Introduction

Alcohol use disorder (AUD) is a significant public health concern, with ∼10% of adults in the United States diagnosed in the past year^1^. Treatment options for AUD remain limited, with only three FDA-approved medications: disulfiram, naltrexone, and acamprosate. The efficacy of these treatments is generally regarded as low to moderate, highlighting the need for new therapeutic approaches. The complexity of alcohol as a drug begins with its dualistic effects, ranging from stimulant outcomes such as euphoria, anxiolysis, and social facilitation, to depressant effects like dysphoria, sedation, motor impairment, and slowed reflexes^2,3^. These opposing effects depend on several factors, including dosage and individual variability^4^. Crucially, this duality may play a role in susceptibility to AUD, as research has shown that individuals who experience a stronger stimulant response are more likely to develop alcohol dependence^5,6^. This dual action property of ethanol has been observed in both humans and rodent models^7,8^. This behavioral complexity may stem from ethanol’s unique chemical structure. Ethanol is a small, two-carbon molecule considerably smaller (46 g/mol) than most other commonly abused drugs, which typically range from 100 to 400 g/mol. Ethanol’s small size contributes to its low potency, low specificity, and rapid, widespread distribution throughout the body, affecting a wide range of tissues and cell types. Unlike most drugs of abuse, ethanol has numerous potential targets with actions on both excitatory and inhibitory neurons^3^. Ethanol potentiates inhibitory receptors like GABA_A_ and glycine receptors, while inhibiting excitatory receptors such as NMDA, AMPA, and metabotropic glutamate receptors (mGluRs)^3^. Additionally, ethanol alters the activity of certain potassium channels, including big potassium (BK) channels, and directly activates G-protein-activated inwardly rectifying potassium (GIRK) channels^9–11^, the focus of this study.

GIRK channels are members of the inwardly rectifying potassium channel family, whereby the inward potassium current is larger than the outward current^12^. However, it is the small outward potassium current that hyperpolarizes neurons, leading to reduced neuronal excitability^12^. GIRK channels are downstream effectors of G protein-coupled receptors (GPCRs) that couple to pertussis toxin-sensitive heterotrimeric G proteins (Gα_i/o_ subfamily), such as the GABA_B_ receptor^13–15^. Interestingly, Baclofen, an agonist for the GABA_B_ receptor, is being prescribed to treat AUD in France^16^ and is being assessed in clinical trials for off-label use in the US^17^. Baclofen leads to activation of GIRK channels via G protein Gβγ subunits^13,18,19^. We reasoned that targeting activation of GIRK channels, which are downstream of GABA_B_ receptors, might provide an alternative approach to treating AUD but with fewer side effects than Baclofen. Another rationale for targeting GIRK channels for treatment of AUD is supported by prior studies that show activation of GIRK channels opposes some of the addictive effects of ethanol. In mice, genetic deletion of GIRK2 increases ethanol self-administration compared to wild-type mice^20^. Reduced expression of GIRK3 has also been implicated in ethanol-related behaviors, with increased binge-like drinking^21^ and ethanol-conditioned place preference^22^. In studies with human neurons, up-regulation of GIRK2 is neuroprotective to the cellular and metabolic effects of ethanol treatment^23,24^.

To test the therapeutic potential of directly activating GIRK channels, we examined the effects of a recently described GIRK channel activator, GiGA1^25^, in ethanol-related behaviors in mice. GiGA1 selectively and potently activates GIRK channels and has rapid and good brain bioavailability in DMPK studies^25,26^. GiGA1 is selective for activating GIRK1/GIRK2 heterotetramers, which are the most common in the brain^12,27^. In contrast to ethanol’s mM affinity for GIRK channels, GiGA1 is effective at µM concentrations^25^. Lastly, GiGA1 administered systemically leads to rapid changes in brain excitability in mice, with recent studies demonstrating anticonvulsant properties in two different mouse epilepsy models^25,28^. Here, we examined the effects of systemic administration of GiGA1, which is applicable to potential future studies with humans, on two different ethanol-dependent behaviors in both male and female mice. We also compared the effects of systemic Baclofen with that of GiGA1, which revealed important advantages of targeting GIRK channels directly with GiGA1. Lastly, we conducted a brain-wide activity mapping using c-Fos expression to reveal potential brain regions where GiGA1 may counteract the effects of ethanol.

## Results

### GiGA1 pre-treatment disrupts ethanol-induced CPP and locomotor stimulation

CPP is a Pavlovian behavioral assay used to assess the motivational and contextual memory effects of rewarding drugs^29^. Whereas CPP with psychostimulants is robust and consistent^30^, prior CPP studies with ethanol have had mixed results^29^. We therefore tested two protocols that focus on dose, and the length and frequency of conditioning trials (**Fig. 1A**). Protocol A used a low dose of ethanol (1.2 g/kg) and short conditioning trials (two 5-min CS+/− sessions)^31,32^. Protocol B, on the other hand, used a dose of 2 g/kg ethanol and five 30-min CS+/− sessions^33^. The two protocols were designed to disentangle the stimulating/rewarding properties from the sedative/aversive properties of ethanol^34^. Also, our experimental design used separate male and female groups (age: 8–12 weeks) since C57BL/6J are known to exhibit sex-differences for alcohol-related behaviors^35^. Ethanol-dependent CPP was assessed by comparing the time spent on the Paired-side before (Pre-test) and after conditioning (Test-day); a positive statistical difference indicated Ethanol-CPP (**Fig. 1B,C**). Ethanol-CPP was observed in females using both protocols (n=12; Protocol A: p=0.0122, Protocol B: p=0.0006, Fig**. 1Di**; see Supplemental Table 1 for details on statistics). For males, we observed ethanol-CPP in a larger Cohort of 22 males (Protocol A: p=0.0360, Protocol B: p=0.0012, **Fig. 1Dii**). A smaller Cohort of twelve males did not reach significance (see Supplemental Table 1). Both protocols appeared to produce ethanol-CPP equally well across sexes when adequately powered (see Supplemental Table 1). We next examined blood alcohol concentration (BAC) with the two protocols. The 2 g/kg of ethanol in Protocol B produced a rapid increase in BAC, exceeding 0.30 g/dL (equivalent to severe intoxication in humans) within 5 minutes and lasting 30–45 minutes (**Fig. 1E**). Conversely, the 1.2 g/kg ethanol (Protocol A) resulted in lower BAC levels, reaching 0.15 g/dL in males and 0.20 g/dL in females (**Fig. 1E**), which are comparable to those reported in Drinking in the Dark (DID) paradigms, where mice typically achieve blood ethanol concentrations exceeding 0.10 g/dL within a 2- to 4-hour access window^36^. Locomotor responses also varied across conditions. At 1.2 g/kg, females exhibited ethanol-induced hyperactivity, a stimulatory effect associated with an increased likelihood of ethanol consumption in both humans and rodents^7,8^ (**Fig. 1F,H**). This effect was absent in males (**Fig. Fii**). At 2 g/kg, males showed locomotor depression from the beginning of conditioning **(Fig. 1Fii**), while in both sexes showed locomotor depression in the final 5 minutes of the trial (**Fig. 1G, I**). Based on the lower BAC and reduced depressant effects with the lower concentration of ethanol, we used Protocol A for subsequent experiments.

**Figure 1.**
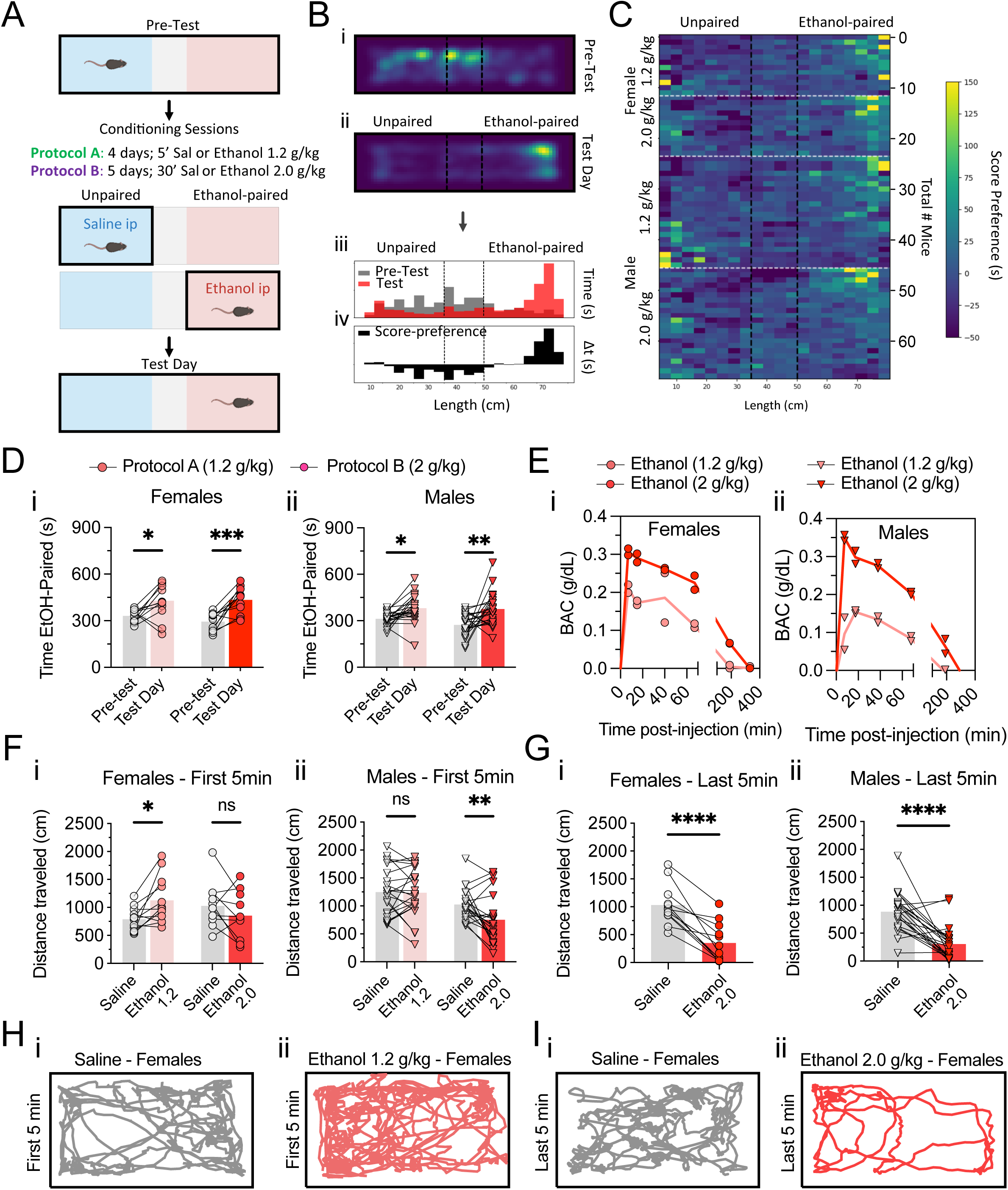
Comparison of ethanol-CPP in female and male mice with two different conditioning protocols. (**A**) Schematic of the ethanol-CPP experimental design. Mice underwent a pre-test session to assess baseline preference, followed by conditioning sessions pairing ethanol with one chamber, and a Test-day session to evaluate conditioned preference. **Protocol A** is 1.2 g/kg ethanol with two 5-min sessions. **Protocol B** is 2.0 g/kg ethanol with five 30-min sessions. (**B**) Heatmaps show spatial occupancy for a representative mouse during the pre-test (**Bi**) and test day (**Bii**) sessions. Increased time spent in the ethanol-paired side is evident after conditioning. Below, histograms quantify time spent across the chamber length (cm) during pre-test (gray) and test day (red) (**Biii)**, with a difference histogram indicating preference score (black) (**Biv**). (C) Heatmap visualization shows preference scores across the chamber length for individual mice, with each row representing one mouse. (**D**) Time spent in the ethanol-paired side during pre-test and test day in females (**Di**, circles) and males (**Dii**, triangles) using Protocol A (light red) or Protocol B (dark red). (**E**) Time-course of blood alcohol concentration (BAC) in females (**Ei,** circles) and males (**Eii,** triangles) after ethanol administration of 1.2 g/kg (light red) or 2.0 g/kg (dark red), corresponding to doses used in Protocols A and B, respectively. (**F**) Distance traveled during the first 5 minutes of conditioning for females (**Fi**) and males (**Fii**) treated with 1.2 g/kg (light red) or 2.0 g/kg (dark red) ethanol. (**G**) Distance traveled during the last 5 minutes of conditioning sessions in Protocol B only, for females (**Gi**) and males (**Gii**). (**H,I**) Representative track maps showing locomotor trajectories of two female mice during conditioning: saline (**Hi**) vs. ethanol (1.2 g/kg) (**Hii**) for the first 5 min, and saline (**Iii**) vs. ethanol (2.0 g/kg) (**Iii**) for the last 5 min. N = 34 mice per protocol (12 females, 22 males). Data are presented as individual mice with mean (bar). Main effects of session (pre-test vs. test day), sex, and their interaction were assessed using two-way ANOVA, with Šídák’s multiple comparisons test used for post hoc analysis and paired t-tests compare saline vs. ethanol treatment within subjects. *p < 0.05, **p < 0.01, ***p < 0.001, ****p < 0.0001. Schematic created with BioRender.com (A).

We next investigated the effects of GiGA1, which selectively activates GIRK1/GIRK2 channels^25^, on ethanol-related behaviors (**Fig. 2A**). Mice were pretreated with GiGA1 (30 mg/kg) 30 minutes before ethanol conditioning trials using Protocol A (**Fig. 2B**). We chose 30 mg/kg because this dose showed clear central nervous system effects previously^25^. As above, both male and female mice exhibited significant ethanol-CPP (**Fig. 2C-D**). Importantly, GiGA1 alone did not produce CPP or conditioned place aversion (CPA). Strikingly, injecting GiGA1 30 minutes prior to ethanol CS+ conditioning trials prevented the expression of ethanol-CPP in both sexes (**Fig. 2C**). GiGA1 reduced locomotion during conditioning sessions in both males and females, consistent with previous findings^25^ (**Fig. 2E-F**). In females, pre-treatment with GiGA1 also prevented the stimulatory effect of ethanol (**Fig. 2E,G**). To determine whether reduced locomotor activity with GiGA1 might indirectly affect conditioning, we examined the effect of GiGA1 on cocaine-CPP. Using Protocol A but with 15 mg/kg cocaine i.p. injections, we examined the effect of GiGA1 (30 mg/kg) on cocaine-CPP. In contrast to ethanol-CPP, GiGA1 did not interfere with acquisition of cocaine-CPP (Supplemental Figure S1 A). To determine if systemic GiGA1 affects memory-based tasks, we also investigated the effect of GiGA1 in the novel object recognition (NOR) task^37^. GiGA1 administered 30 minutes before object exploration (acquisition phase) did not impair novel object recognition memory (Fig. S1 B). Taken together, these results indicate that systemic GiGA1 pre-treatment during conditioning appears to selectively prevent the acquisition of ethanol-CPP in both males and females.

**Figure 2.**
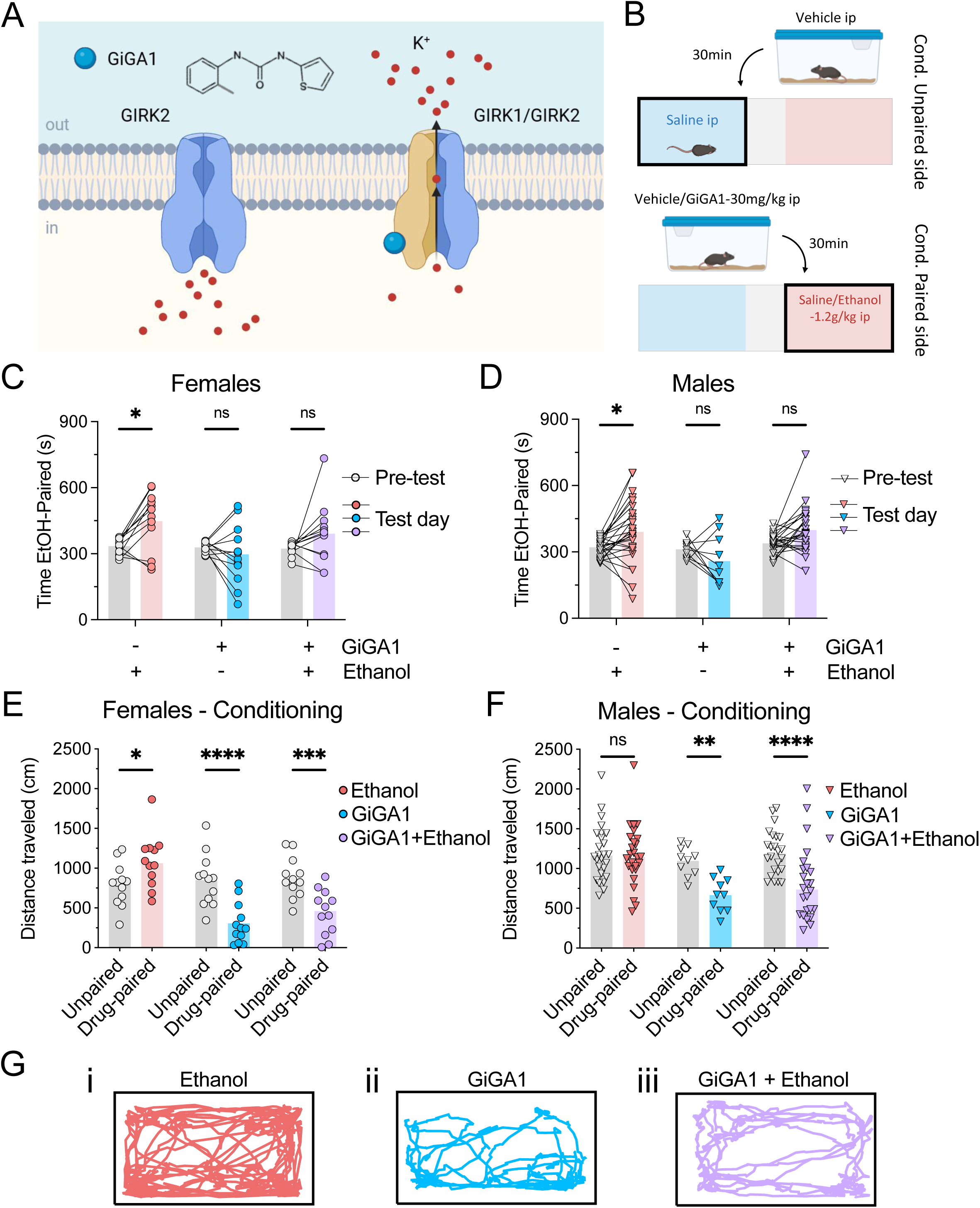
GiGA1 occludes ethanol-CPP acquisition in both female and male mice. (**A**) Chemical structure of GiGA1 and schematic cartoon illustrating its selective activation of GIRK1/GIRK2 channels, leading to outward potassium efflux^25^. (**B**) Experimental timeline for GiGA1 intervention. Mice received GiGA1 (30 mg/kg, i.p.) 30 minutes prior to each conditioning session using Protocol A (1.2 g/kg ethanol, two 5-min sessions). (**C-D**) Time spent in the ethanol-paired side during Pre-test and Test Day sessions in females (**C**, circles) and males (**D**, triangles), across three treatment groups: ethanol alone (light red), GiGA1 alone (blue), and GiGA1–ethanol co-treatment (purple). GiGA1 administration prevented the development of ethanol-induced CPP in both sexes. (**E-F**) Locomotor activity during conditioning sessions, measured as total distance traveled, in females (**E**) and males (**F**) across the same three treatment groups. (**G**) Representative track maps showing locomotor trajectories during conditioning sessions under different treatments: (**Gi**) ethanol, (**Gii**) GiGA1, and (**Giii**) GiGA1 plus ethanol. N = 96 mice (36 females, 60 males). Data are presented as individual mice with mean (bar). Main effects of session (pre-test vs. Test-day), treatment (Vehicle vs. GiGA1), and their interaction were assessed using two-way ANOVA, with Šídák’s multiple comparisons test used for post hoc analysis. *p < 0.05, **p < 0.01, ***p < 0.001, ****p < 0.0001. Schematic created with BioRender.com (**A** and **B**).

### GiGA1 decreases ethanol voluntary drinking without affecting water consumption

Having shown that GiGA1 prevents acquisition of ethanol-dependent CPP, we next investigated the effect of GiGA1 on ethanol consumption in a self-administration paradigm. In this experiment, we investigated whether GiGA1 could reduce ethanol drinking in mice that had already developed high preference and intake of ethanol, more akin to human consumption. We implemented a mixed model combining two-bottle choice (2BC) with the drinking-in-the-dark (DID) paradigm (2BC/DID)^38,39^ (see **Fig. 3A**, Cohort 1, and Methods for details). Male and female C57BL/6J mice, a strain known for high ethanol consumption^40^, were given limited access to ethanol for six weeks under a two-bottle choice paradigm to support binge drinking (**Fig. 3A**). Intake stabilized by week four at 2.3 g/kg/2h on average (**Fig. 3B, red line**), consistent with previous binge-drinking models^36^. Females consumed significantly more ethanol than males in the latter weeks of the protocol (**Fig. 3B**, see Supplemental Table 1). Ethanol preference remained high (∼90%) from week two onward and indistinguishable between males and females (**Fig. 3C**). We quantified ethanol consumption in week seven using a lickometer, which provided high-resolution, real-time measurement of drinking behavior (**Fig. 3A**). Both licking behavior and ethanol intake (measured by volume consumed) were assessed, with each mouse tested under vehicle and GiGA1 conditions (**Fig. 3D,E**). In Cohort 1, injection of GiGA1 (30 mg/kg) 15 minutes before a session significantly reduced the number of licks and ethanol intake (**Fig. 3E-G**). A significant linear relationship between ethanol intake and the number of ethanol licks was observed for both the vehicle (R² = 0.40, n = 15, p < 0.05) and GiGA1 groups (R² = 0.26, n = 15, p < 0.05) (**Fig. 3E**). By contrast, injection of GiGA1 had no effect on water consumption (**Fig. 3H,I**). Although mean ethanol preference was lower following GiGA1 treatment (55 ± 5) compared to vehicle (68 ± 5), the difference was not statistically significant in this Cohort (unpaired t-test, p = 0.1159; **Fig. 3J**). Dose-response experiments testing GiGA1 at 20, 30, and 40 mg/kg revealed that doses >20 mg/kg significantly reduced ethanol intake (**Fig. S2**). Importantly, GiGA1 did not affect 4% sucrose consumption at the end of the experiment (**Fig. 3K**), indicating a selective effect on ethanol intake. Overall, these findings show that activating GIRK1/GIRK2 channels with GiGA1 significantly reduces voluntary ethanol consumption, without affecting drinking behavior (i.e. water) or consumption of natural rewards (i.e., sucrose).

**Figure 3.**
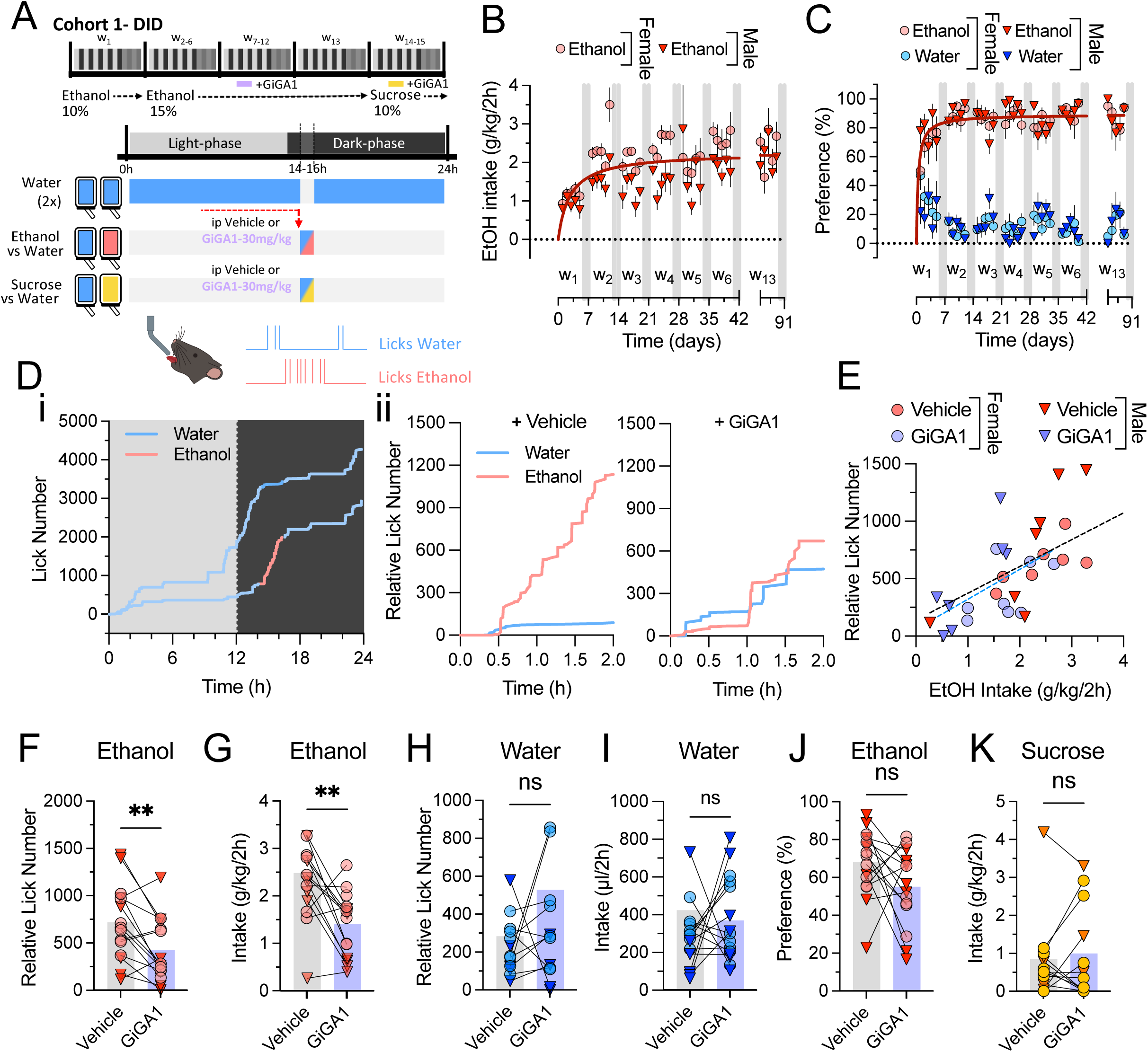
GiGA1 reduces voluntary ethanol consumption while preserving water and sucrose intake in a 2BC-DID lickometer paradigm. (**A**) Schematic of the 2BC-DID experimental design. Female and male mice (Cohort 1) were tested over 15 weeks. Single-housed mice had 2 h/day access (Mon–Fri, 2 h into the dark cycle) to 10% ethanol in week 1 and 15% ethanol thereafter. GiGA1 (30 mg/kg, i.p.) effects were measured in a lickometer cage: mice underwent 14 h habituation, received GiGA1 or vehicle i.p. (purple bar), and then completed the usual 2 h drinking session. At the end of EtOH administration, mice were exposed to 4% sucrose and tested again with GiGA1 (yellow bar). (**B**) Ethanol intake (red) and (**C**) ethanol preference (red) versus water (blue) across the duration of the experiment for females (circles) and males (triangles). (**D**) Lick number plotted over time for a representative mouse: (**Di**) 24-hour time-course including habituation, drinking, and post-session periods. (**Dii**) Expanded views of the 2-hour drinking session with vehicle (left plot) and GiGA1 (right plot) for the same mouse. (**E**) Plot of ethanol licks (number) versus ethanol intake (bottle weight) for vehicle and GiGA1 sessions (each symbol = one mouse/session). Line shows positive correlation. (**F,G**) Plot of the relative lick number and ethanol intake (red) for females (circles) and males (triangles) with vehicle (grey bar) or GiGA1 (purple bar) treatment. (**H,I**) Plot of the relative lick number and water intake (blue) for females (circles) and males (triangles) with vehicle (grey bar) or GiGA1 (purple bar) treatment. (**J**) Plot of drinking preference for ethanol as a proportion of total fluid intake. (**K**) Plot of sucrose (4%, yellow symbols) intake with vehicle or GiGA1 treatment in same Cohort (see Methods for details). N = 15 mice (8 females shown as circles, 7 males as triangles). Data are presented as individual mice with mean (bar). Paired t-tests were used to compare vehicle vs. GiGA1 treatment in **F–K**. **p < 0.01. Schematic created with BioRender.com (**A**).

To investigate potential sex differences and also measure BAC, male and female C57BL/6J mice underwent the same ethanol-2BC/DID protocol but remained in their home cage (**Fig. 4A**, Cohort 2). Ethanol consumption increased in both sexes over time, with females again exhibiting significantly higher intake levels than males starting from week 2 (**Fig. 4Bi**), as shown previously^41,42^. No significant sex differences were observed in ethanol preference, reaching approximately 90% preference by week 4 (**Fig. 4Bii**). After four weeks of stable ethanol consumption, we administered either vehicle or GiGA1 (i.p.) 30 minutes before the drinking sessions. In addition to measuring volume of consumption, we also measured BAC in blood draws (**Fig. 4A**). In Cohort 2, GiGA1 significantly reduced ethanol intake in both male and female mice without affecting water consumption (**Fig. 4C,D**). GiGA1 treatment significantly reduced ethanol preference in females but not in males (**Fig. 4E,F**). Interestingly, GiGA1 significantly decreased BAC in both sexes, with BAC levels dropping below 0.05 g/dL (**Fig. 4G-J**). After the ethanol DID paradigm, water bottles were replaced with water/sucrose in their home cages and mice were allowed to habituate to sucrose for one week before testing GiGA1. In Cohort 2, GiGA1 appeared to reduce sucrose intake in both males and females (**Fig. 4K,L**). This effect of GiGA1 may be influenced by the prior one week habituation period (**Fig. 4A**), which differed from Cohort 1 (see Discussion). Importantly, GiGA1 significantly reduced ethanol consumption without affecting water intake in both Cohorts.

**Figure 4.**
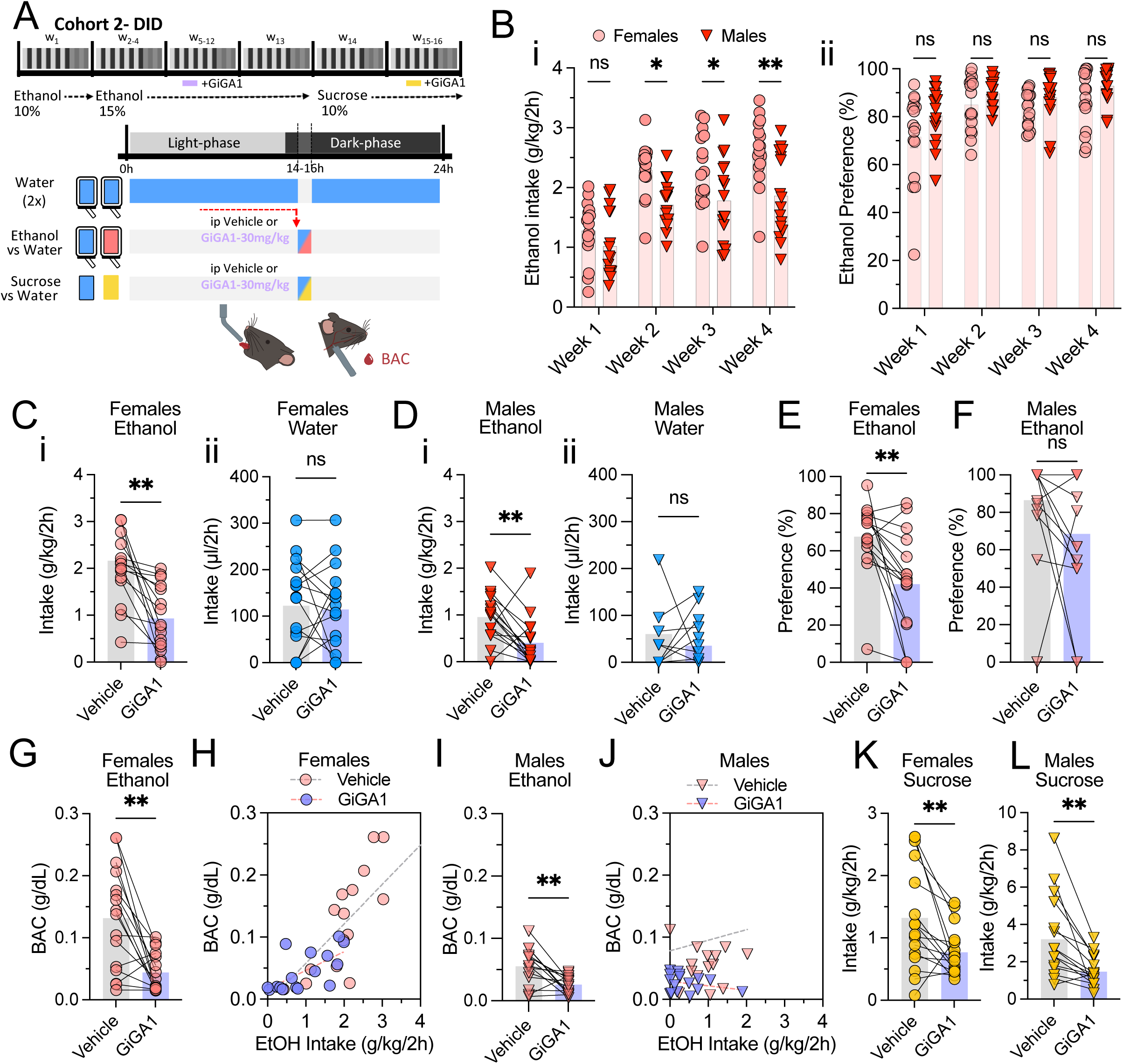
GiGA1 reduces ethanol intake and BAC in both female and male mice in a 2BC-DID home cage paradigm. (**A**) Schematic of the 16-week 2BC-DID protocol for Cohort 2, where female and male mice were tested separately under the same regimen as Cohort 1 (see Methods for details). The only modifications were that drinking sessions took place in standard home cages instead of lickometer cages, BAC measurements were added at the end of each GiGA1 (30 mg/kg, i.p.) or vehicle session, and mice were exposed to 4% sucrose for 1 week, and then tested again with GiGA1 (yellow bar). (**Bi**) Baseline ethanol intake and (**Bii**) ethanol preference during the 4 weeks preceding treatment, showing statistically higher consumption in females. (**C–F**) Plots show the effect of GiGA1 on ethanol (red) and water (blue) intake in females (**C,E, circles**) and males (**D,F, triangles**). (**E,F**) GiGA1 effects on ethanol preference for females and males. (**G-J**) Plots show the effect of GiGA1 effects on BAC in females (**G,H**) and males (**I,J**). Positive correlation between ethanol intake and BAC in the same female (**H**) and male (**J**) mice. (**K,L**) Plots show the effect of GiGA1 on Sucrose (4%) intake with vehicle or GiGA1 treatment in the same Cohort, one week after habituation to sucrose consumption. N = 32 mice (16 females, circles; 16 males, triangles). Data are presented as individual mice with mean (bar). Main effects of session (Week number), sex, and their interaction were assessed using two-way ANOVA, with Šídák’s multiple comparisons test used for post hoc analysis and paired t-tests were used to compare vehicle vs. GiGA1 treatment in **D–L**. *p < 0.05, **p < 0.01. Schematic created with BioRender.com (**A**).

### Baclofen does not block ethanol-CPP but reduces ethanol’s stimulatory effect and drinking

Baclofen, which activates GIRK channels through GABA_B_ receptors coupled to G proteins (**Fig. 5A**), is a potential treatment for AUD^43–45^. We first investigated the effect of Baclofen on ethanol-CPP, to compare with GiGA1 (**Fig. 5B**). We first focused on female mice because of their more robust ethanol-CPP, stronger stimulatory responses, and higher ethanol intake, compared to males. We tested 7.5 mg/kg, a dose used previously for CPP^46^, and found that Baclofen did not block acquisition of ethanol-CPP (**Fig. 5C**). Interestingly, while Baclofen did not affect acquisition of ethanol-CPP, it did reduce ethanol-induced hyperactivity observed during the acquisition phase, like GiGA1 (**Fig. 5D,E**). Additionally, Baclofen reduced locomotor activity in the absence of ethanol (**Fig. 5D,E**). Thus both GiGA1 and baclofen exhibited some properties of sedation, though this did not interfere with acquiring ethanol dependent CPP after Baclofen administration.

**Figure 5.**
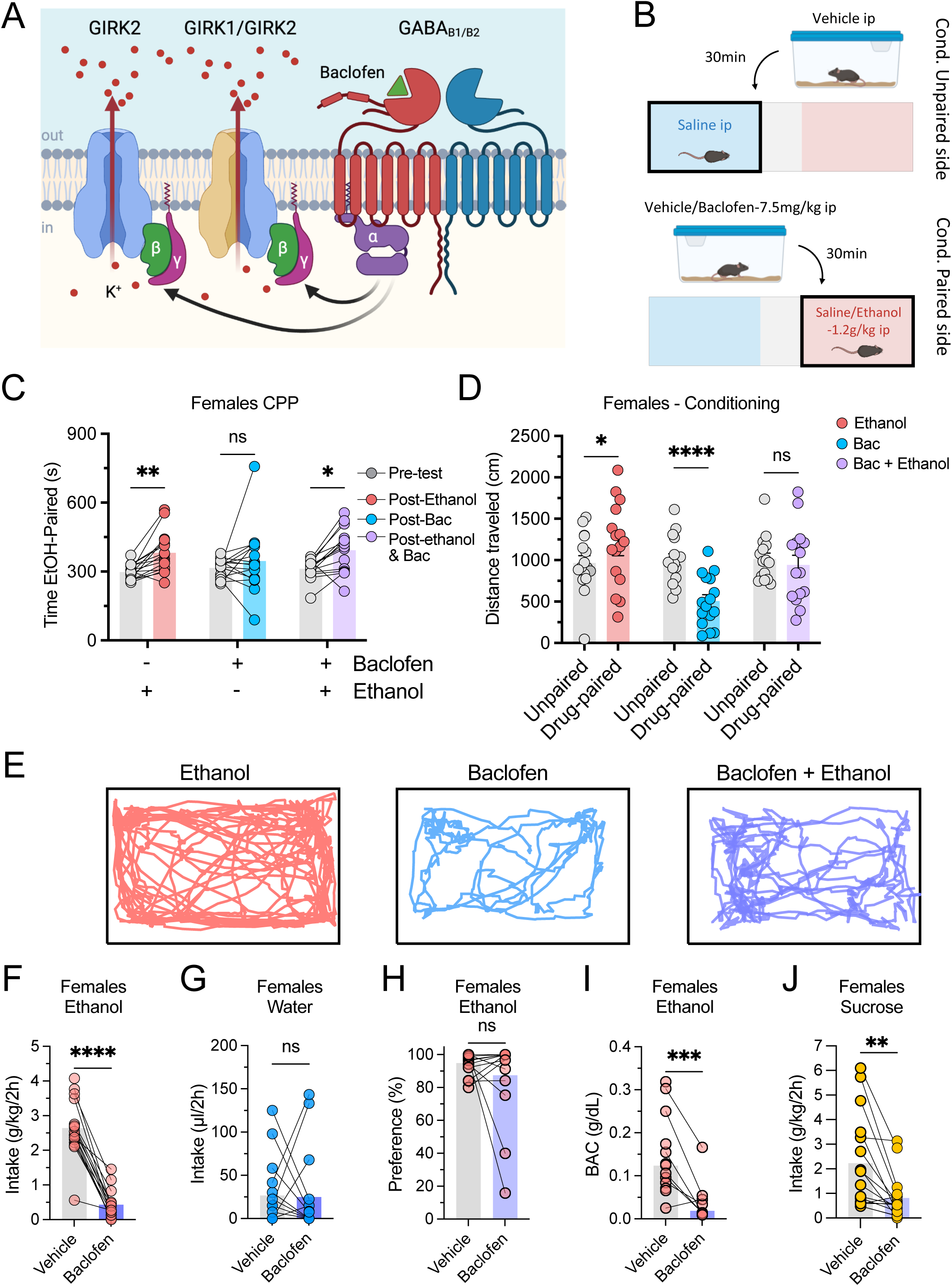
Baclofen reduces voluntary ethanol intake but does not occlude ethanol-CPP. (**A**) Schematic cartoon illustrates Baclofen-induced GIRK channel opening via GABA_B_ receptor activation and Gβγ subunit signaling, leading to outward potassium efflux. (**B**) Experimental timeline for Baclofen intervention. Female mice received Baclofen (7.5 mg/kg, i.p.) or vehicle 30 minutes prior to each ethanol conditioning session (Protocol A: 1.2 g/kg ethanol, two 5-min sessions; see Methods). (**C**) Time spent in the ethanol-paired side during pre-test and test day sessions in females, across three treatment groups: ethanol alone (red), Baclofen alone (blue), and Baclofen–ethanol co-treatment (purple). Baclofen did not prevent ethanol-CPP. (**D**) Locomotor activity during conditioning sessions, measured as total distance traveled, in females across the same three treatment groups. (**E**) Representative track maps show locomotor trajectories during conditioning sessions under different treatments: (left) ethanol, (middle) Baclofen, and (right) Baclofen plus ethanol. (**F,G**) Plots show ethanol (red) and water (blue) intake for vehicle (grey bar) and Baclofen (purple bar) treatments. (**H**) Plot shows drinking preference for ethanol vs water. (**I**) BAC measurements for vehicle and Baclofen treatments following ethanol intake. (**J**) Plot shows Sucrose (4%) intake with vehicle (grey bar) or Baclofen (purple bar) treatment, following one week of habituation to sucrose consumption, using same drinking regime as for ethanol. Similar results were obtained in males (see **Supplemental Fig S2**). N = 47 female mice (15–16 per group) for Ethanol-CPP experiments (panels **C–D**). Main effects of session (pre-test vs. Test day), treatment (Vehicle vs. Baclofen), and their interaction were assessed by two-way ANOVA with Šídák’s multiple comparisons for post hoc testing. N = 16 females (circles) were used for the 2BC-DID paradigm (**F–J**). Paired t-tests compared vehicle vs. Baclofen treatments. Data are presented as individual mice with mean (bar). *p < 0.05, **p < 0.01, ***p < 0.001, ****p < 0.0001. Schematic created with BioRender.com (A and B).

We next examined the effect of Baclofen on voluntary ethanol consumption. Similar to GiGA1, Baclofen significantly reduced ethanol intake and lowered BAC (**Fig. 5F,I**) without affecting water consumption (**Fig. 5G**). In contrast to GiGA1, however. Baclofen did not significantly reduce ethanol preference in females (**Fig. 5H**). Baclofen also reduced sucrose intake (**Fig. 5J**), which has been shown previously^47,48^ and enhance ethanol sedation in C57BL/6J mice}. These changes with Baclofen were similar with male mice (**Supplemental Fig. S2**). Overall, it seems that systemic Baclofen can reduce ethanol-induced hyperactivity and voluntary ethanol consumption, but does not appear to alter ethanol’s reinforcing properties (i.e., did not block acquisition of ethanol-CPP) or significantly reduce ethanol preference over water.

### GiGA1 blunts ethanol-induced c-Fos changes in key brain regions in female mice

To identify potential brain regions affected by systemic GiGA1 and ethanol, we conducted a brain-wide survey of changes in excitability using the immediate early gene (IEG) c-Fos as a proxy for changes in neuronal activity. We focussed on females because of the more robust CPP, higher ethanol BACs, and paucity of c-Fos studies with females^49–53^. Mice were injected with either GiGA1 (30 mg/kg) or vehicle and allowed to rest in their home cage for 30 minutes; mice were then injected with ethanol (1.2 g/kg) or saline and placed in a novel open-field environment for 60 minutes before being sacrificed (**Fig. 6A**). We then quantified the c-Fos+ neurons per mm^2^ across the four experimental groups in specific brain regions (**Fig. 7A-L**). We also calculated fold-change relative to saline (**Fig. 7M**). We hypothesized that GiGA1 pretreatment would prevent ethanol-induced changes in c-Fos expression in brain regions where GIRK1 is expressed (**Fig. 6A,B**). We first examined the effect of ethanol alone in females. At 60 minutes following a single i.p. injection of 1.2 g/kg ethanol (**Fig. 6A**), c-Fos expression increased in several regions compared to saline i.p. injection, including the rostral part of the piriform cortex (Pir), the paraventricular nucleus of the thalamus (PVT), the paraventricular nucleus of the hypothalamus (PVN) and the Edinger-Westphal nucleus (EW), consistent with the literature^54^ (**Figs. 6B, 7A, 7J, 7K, 7M**). Importantly, GiGA1 pre-treatment attenuated the ethanol-induced c-Fos increase in the Pir, PVT, PVN and EW (**Fig. 6B, 7A** [Pir], **7J** [PVT], **7K** [PVN], **7L** [EW]). In some brain regions, ethanol caused a significant reduction in c-Fos expression across hippocampal regions (CA1-3 and DG) and in retrosplenial granular/dysgranular (RSGc/RSD) cortices (**Figs. 6B & 7C**), consistent with prior studies showing that novelty-induced c-Fos activity is suppressed by ethanol^55,56^. Similarly, GiGA1 administered alone or in combination with ethanol also produced a significant reduction in these regions (**Fig. 7F** [DG], **7G** [CA3], **7C** [RSGc], and **7M**). This finding is noteworthy given the high expression of GIRK1 in the hippocampus and retrosplenial cortex. Additionally, in the dorsal lateral striatum (DLS), ethanol also lowered c-Fos, consistent with earlier findings at this dose in the C57BL/6J strain^57^. GiGA1, alone or combined with ethanol, similarly reduced c-Fos relative to vehicle (**Fig. 7H** [DLS], and **7M**).We next assessed changes in c-Fos expression in brain regions previously implicated in ethanol exposure, including the amygdala, nucleus accumbens, prefrontal cortex, and midbrain. Ethanol significantly increased c-Fos levels in the central amygdala (CeA), but not in the basomedial amygdala (BMA) or basolateral amygdala (BLA), consistent with previous reports showing ethanol-induced activation restricted to the CeA (**Fig. 6B, 7E** [CeA], **7D** [BLA], and **7M**). In the nucleus accumbens (NAc) core, ethanol increased the mean c-Fos/mm^2^ (though not significantly), while GiGA1 showed an overall main effect to reduce c-Fos expression (**Fig. 6B, 7I** [NAc], and **7M**). A similar pattern was observed in the ventral tenia tecta (VTT) (**Fig. 7B** [VTT], and **7M**). In the prelimbic (PrL) and cingulate (Cg) cortices, no significant differences in c-Fos were detected across treatment groups. The ventral tegmental area (VTA) and substantia nigra (SN) showed uniformly low c-Fos levels. In contrast, the anterior pretectal nucleus (APT) exhibited a robust reduction following GiGA1 treatment, both alone and in combination with ethanol (**Fig. 7M**).

**Figure 6.**
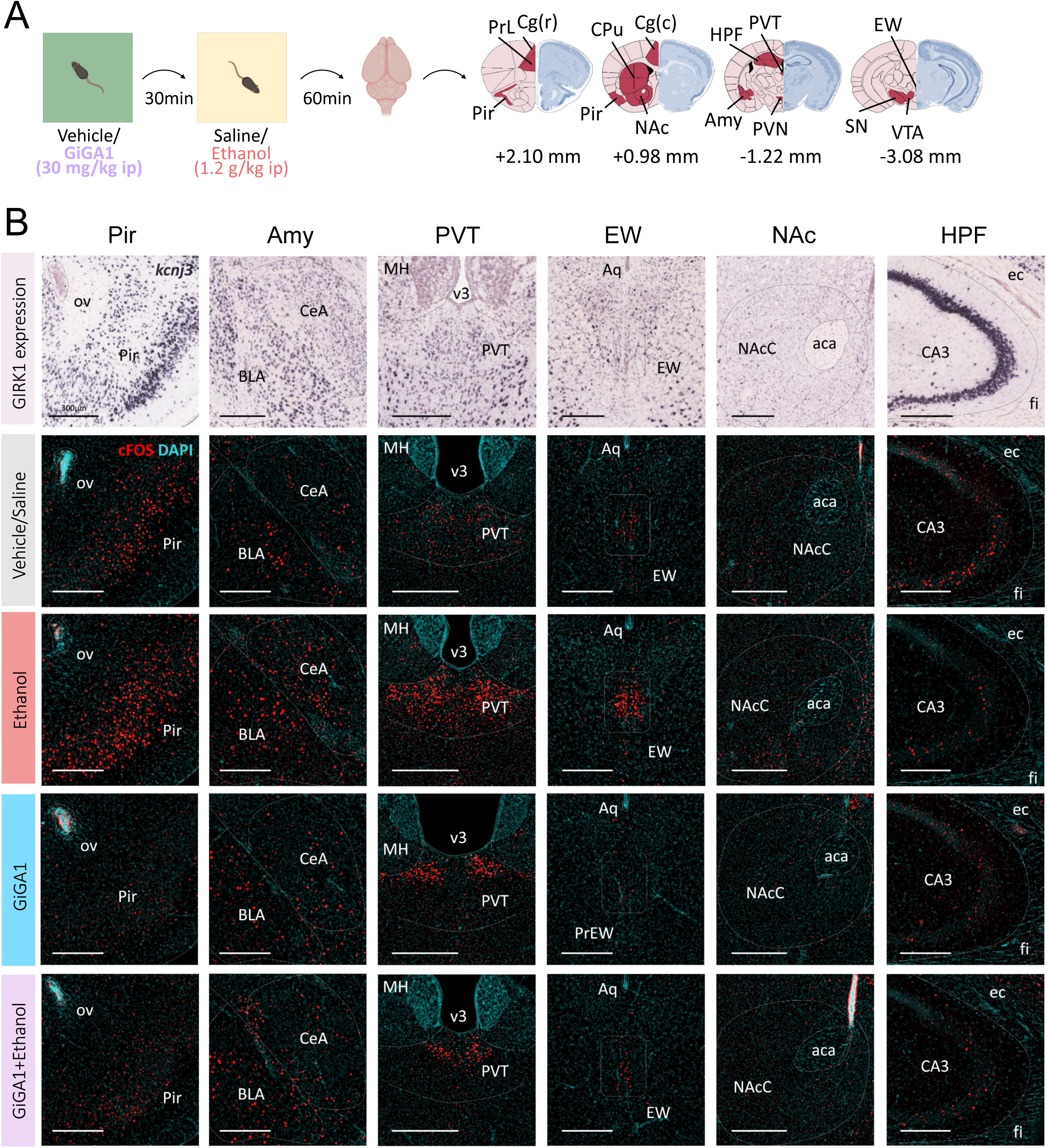
GiGA1 alters ethanol-induced c-Fos activation across multiple brain regions. (**A**) Timeline for c-Fos experiments: mice received vehicle or GiGA1 (30 mg/kg, i.p.), rested for 30 min, and then received saline or ethanol (1.2 g/kg, i.p.). After 1 h, brains were harvested, fixed, coronally sectioned, immunolabeled, and imaged. Four coronal sections per animal were chosen to include regions previously shown to be altered by ethanol^54^ and areas with high GIRK1 expression. Ethanol-sensitive regions shown in red and GIRK1-rich regions shown in dark blue. (**B**) Representative images for six regions—piriform cortex (Pir), basolateral/central amygdala (Amy), paraventricular nucleus of the thalamus (PVT), Edinger–Westphal nucleus (EW), nucleus accumbens core (NAc), and hippocampal formation CA3 (HPF). Top row: *Kcnj3* mRNA in situ hybridization (Allen Mouse Brain Atlas^79^). Rows 2–5: images shown are stained with DAPI (blue) and for c-Fos (red) staining for the indicated conditions. Schematic created with BioRender.com (**A**).

**Figure 7.**
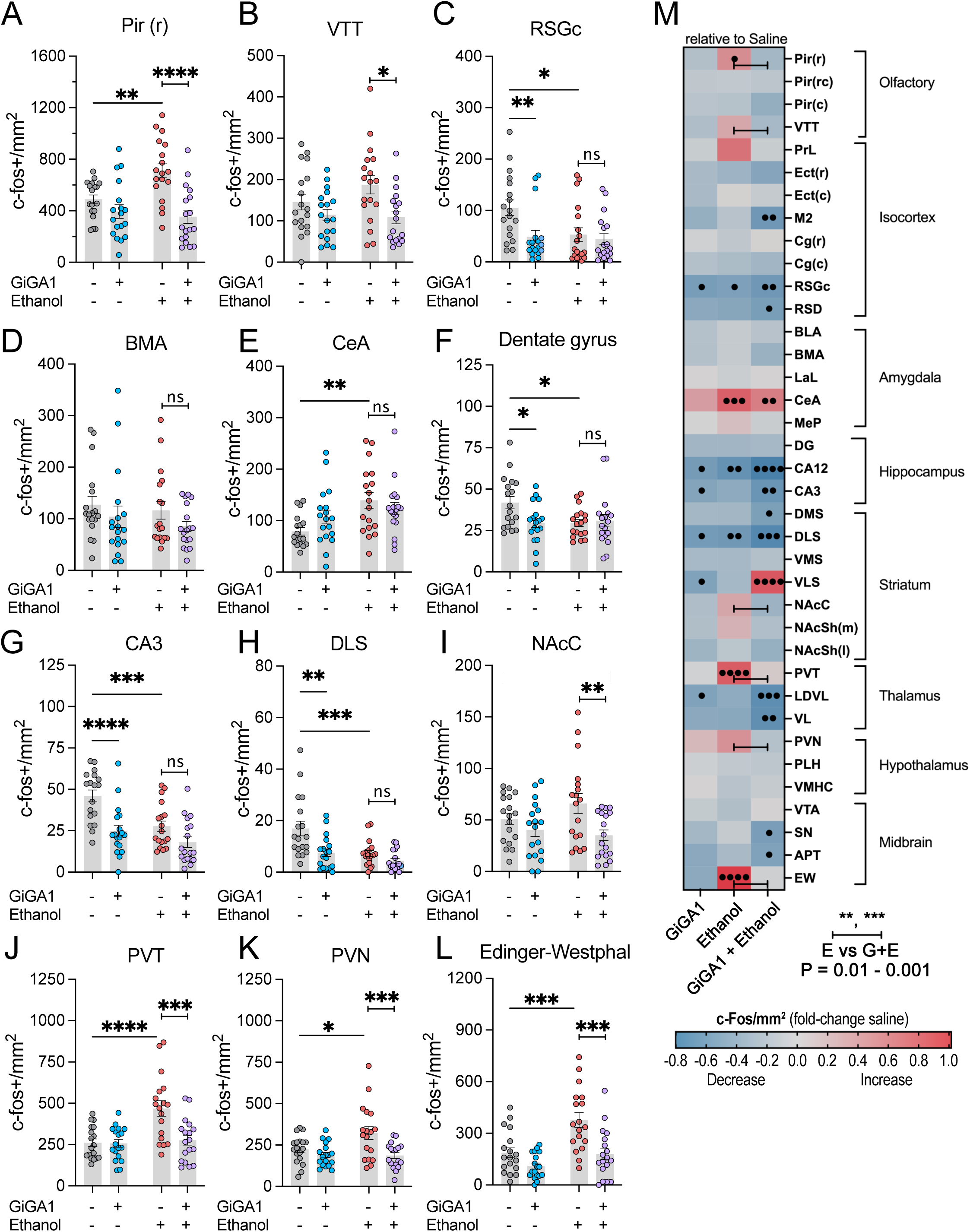
GiGA1 occludes ethanol-induced c-Fos activation in multiple brain regions. (**A–L**) Quantification of c-Fos⁺ cell density (cells/mm^2^) under Control (grey, no GiGA1 or ethanol), ethanol (red), GiGA1 (blue), and GiGA1–ethanol (purple) conditions in (**A**) Pir (r), (**B**) ventral tenia tecta (VTT), (**C**) retrosplenial granular cortex, (**D**) basomedial amygdala, (**E**) central amygdala (CeA), (**F**) dentate gyrus, (**G**) CA3 field, (**H**) dorsolateral striatum (DLS), **(I**) NAc core (NacC), (**J**) paraventricular nucleus of the thalamus (PVT), (**K**) paraventricular nucleus of the hypothalamus (PVN), and (**L**) Edinger–Westphal (EW) nucleus. (**M**) Heatmap shows fold-change in c-Fos⁺ density (relative to saline) across 37 regions, grouped into olfactory, isocortex, amygdala, hippocampus, striatum, thalamus, hypothalamus, and midbrain. Red = increased expression; blue = decreased expression; gray = no change. N = 36 mice (9 per group). In panels **A–L**, main effects of ethanol, GiGA1, and their interaction were assessed by two-way ANOVA with Šídák’s post hoc tests. The Šídák’s post hoc comparisons is indicated by asterisks (*p < 0.05; **p < 0.01; ***p < 0.001; ****p < 0.0001). Significant main effects of GiGA1 are indicated with hash symbols (#p < 0.05; ##p < 0.01). All data are mean and individual mice. (**M**) heatmap of the fold change in c-Fos relative to saline, for ethanol alone, GiGA1 alone and GiGA1 plus ethanol. Bar shows significant change between Ethanol and Ethanol+GiGA1 from (A-L). (• p < 0.05; • • p < 0.01; • • • p < 0.001; • • • • p < 0.0001)

Taken together, our brain-wide c-Fos study showed that GiGA1 reduced basal activity in GIRK1-rich hippocampal and cortical areas, thereby mimicking the effect of ethanol in these areas, but significantly attenuated ethanol-evoked activation in key reward and stress centers (e.g., Pir, PVT, PVN, EW), supporting a mechanism in which GIRK1/GIRK2 channel activation counteracts ethanol’s circuit-level actions in those areas.

## Discussion

Here, we demonstrate for the first time that the small molecule GiGA1, a selective GIRK1/GIRK2 channel activator, can significantly reduce alcohol-mediated behaviors in two distinct preclinical models of alcohol binge exposure. GiGA1 prevents acquisition of ethanol-CPP in both males and female mice. Importantly, GiGA1 significantly reduces ethanol self-administration without affecting water consumption in a voluntary drinking model (2BC-DID). Notably, this reduction in ethanol consumption leads to BAC below the intoxication threshold, but does not produce complete abstinence. When compared to Baclofen administration, GiGA1 shows distinct advantages. While Baclofen reduces ethanol intake similarly to GiGA1, Baclofen fails to prevent acquisition of ethanol-CPP, highlighting GiGA1’s broader functional impact on both consumption and conditioned responses to ethanol. Lastly, GiGA1 modulates ethanol-induced neuronal activation patterns, as inferred by changes in c-Fos expression, reversing ethanol-driven increases in multiple brain regions implicated in AUD. Thus, targeting activation of primarily GIRK1/GIRK2 channels, through a compound like GiGA1, could provide a new approach to treating AUD in humans in the future.

### GiGA1 show pre-clinical efficacy in AUD mouse models

GIRK channels have been implicated in addiction, with ethanol known to directly activate them^12,58^. However, while ethanol broadly interacts with multiple targets in the brain, GiGA1 selectively activates GIRK1/GIRK2 channels, and with greater potency than ethanol^25^. While implicated in the actions of ethanol, GIRK channels are emerging as a protein counteracting the effects of ethanol. For example, mice lacking GIRK2 or GIRK3 self-administer more alcohol^20,21^. Overexpression of GIRK2 in human glutamate neurons appears to provide some protection from the effects of ethanol^24^. Our results with in vivo studies using GiGA1 further strengthen the concept that activating GIRK channels is protective in AUD. We speculate that by selectively enhancing GIRK-mediated inhibitory signaling, GiGA1 could restore inhibitory control in key brain regions affected by ethanol, mitigating its pathological effects and potentially restoring homeostasis in AUD-affected brain areas. This targeted modulation of GIRK channels counteracts ethanol-induced disruptions and may also help mitigate ethanol-induced neuroadaptations. As a result, GiGA1 may shift the balance toward neuroprotection, ultimately reducing alcohol-seeking behaviors and providing a promising therapeutic approach for AUD.

While many preclinical studies rely on only voluntary drinking models, we used both CPP and 2BC-DID paradigms to examine the effect of GiGA1 on the behavioral response to ethanol. The ethanol CPP model enabled us to evaluate how GiGA1 disrupts the learned associations between environmental cues and ethanol reinforcement, while the 2BC-DID model provides insights into the ability of GiGA1 to reduce ethanol consumption in mice with extended drinking history. The 2BC-DID model is particularly valuable as it allows the assessment of GiGA1’s impact on mice that have already developed a preference for self-administering ethanol, providing a more realistic representation of the challenge faced in clinical interventions. The ability of GiGA1 to reduce ethanol intake in mice with a drinking habit combined with its preventing context-specific ethanol-dependent memories underscores its potential for further pre-clinical development.

Unlike psychostimulants and opioids which produce strong CPP^30^, prior studies had shown that ethanol-CPP can be variable^29^. To address this, we evaluated two distinct conditioning protocols for CPP: a long duration, high-dose protocol (five 30-min sessions at 2 g/kg ethanol) and a shorter duration, lower-dose protocol (two 5-min sessions at 1.2 g/kg ethanol). Both protocols successfully produced ethanol-CPP in males and females. However, ethanol-CPP was not robustly expressed in males, likely due to high inter-individual variability and perhaps an attenuated response to ethanol in male mice. Once we increased the sample size by doubling the size of the Cohort, we could reveal significant ethanol-CPP. Consistent with prior studies^41,42,59,60^, females in our study demonstrated more robust ethanol-CPP, higher voluntary ethanol consumption, and a unique stimulatory response to ethanol. Despite established sex differences in AUD-related behaviors, GiGA1 effectively reduced ethanol intake and blocked ethanol-CPP in both males and females using the shorter 1.2 g/kg protocol. Notably, GiGA1 specifically reduced ethanol preference in females, as well as attenuating the locomotor stimulatory effects of ethanol, aligning with previous observations of greater female sensitivity to pharmacological interventions targeting ethanol reinforcement^61^. Although sex differences significantly influence ethanol-induced behaviors, our findings suggest that activating GIRK1/GIRK2 channels can modulate ethanol-responsive circuitry across sexes. Interestingly, sex differences have been observed with Baclofen treatment for AUD in clinical studies. In a randomized trial, females showed a significant therapeutic response at a lower dose of Baclofen (30 mg/day), whereas males showed marginal effects at a higher dose (90 mg/day), with females also experiencing higher rates of sedation-related side effects at the higher dose^44^. These findings underscore the importance of considering sex and dose interactions when developing pharmacotherapies.

We compared the effects of GiGA1 directly with Baclofen, which is being assessed for treating AUD in clinical trials in the US^45^. In contrast to GiGA1, Baclofen did not block ethanol-CPP. While GiGA1 selectively activates GIRK1/GIRK2 channels, Baclofen stimulates GABA_B_ receptors, which activate all GIRK channels through G proteins, and in addition affects numerous other down-stream effectors (e.g., Adenylyl cyclase, PKA). Furthermore, GABA_B_ receptors are expressed in regions or neurons that do not express GIRK channels. For example, GIRK1/GIRK2 channels are predominantly found in GABAergic and glutamatergic neurons, whereas Baclofen can also influence dopamine neurons of the SN and VTA, which lack the GIRK1 subunit^12^. In addition, the midbrain contains GABAergic interneurons and glutamatergic inputs^62,63^, further diversifying potential sites of action. In this regard, Baclofen likely engages additional neuronal populations and circuits that are distinct from GiGA1.

For 2BC, we observed that both GiGA1 and Baclofen significantly reduced voluntary ethanol intake, but GiGA1 reduced ethanol preference in female mice. These results highlight that GiGA1 uniquely influences ethanol-associated reinforcement, emphasizing its potential advantage over Baclofen in targeting mechanisms underlying ethanol preference. It will be important to examine in the future the effect of GiGA1 in models of AUD involving chronic use, such as CIE^21^, or operant-dependent administration of ethanol ^64^.

### Other considerations with activation of GIRK channels in vivo

We found that both GiGA1 and Baclofen reduced locomotor activity. This sedative effect of baclofen is well known and is therefore unsurprising that GiGA1 also produces a similar effect.

We considered whether this effect of GiGA1 could affect ethanol-CPP and ethanol self-administration. A confounding effect of sedation on the behavioral tests with ethanol seems unlikely since GiGA1 had no effect on acquisition of cocaine-CPP or interfering with NOR, a memory-based task. Lastly, Baclofen did reduce locomotor activity but had no effect on ethanol-CPP. Thus, the sedative effect does not appear to affect the acquisition of ethanol-CPP. For 2BC, we observed that both GiGA1 and Baclofen significantly reduced voluntary ethanol intake but did not affect water consumption. Also, GiGA1 reduced ethanol preference in female but not male mice in the 2BC paradigm. Taken together, it seems unlikely that decreases in locomotor activity can explain the reduced ethanol self-administration. Future studies developing analogs of GiGA1 that are more potent and selective may avoid some of these sedative properties.

We also investigated the effect of GiGA1 with self-administration of sucrose using the same 2BC-DID paradigm. We used two slightly different protocols. In Cohort 1, mice were given water/sucrose after the ethanol/water sessions ended (2-4 week gap), while in Cohort 2, we allowed the mice to self-administer sucrose for 1 week, similar to the ethanol protocol, before testing GiGA1. GiGA1 administration did not affect sucrose consumption in Cohort 1, but did reduce sucrose consumption in Cohort 2. Thus, it is possible that GiGA1 has little effect on acute drinking, but does reduce more long-term reward-dependent drinking. Overall, while reductions in sucrose intake and locomotor activity may complicate the interpretation, the consistent suppression of ethanol consumption suggests that GiGA1 exerts a specific and beneficial action on alcohol-related behaviors.

### Brain regions involved in GiGA1 actions

To investigate the impact of GiGA1 and ethanol on neuronal activity, we conducted a brain-wide c-Fos mapping study in female mice. In addition, to brain regions commonly involved in alcohol behaviors (Hippocampus, EW, PVT, PVN, CeA), we also observed ethanol-induced increases in c-Fos expression in the piriform cortex and ventral tenia tecta (VTT). In contrast, ethanol reduced c-Fos expression in the dorsolateral striatum (DLS), retrosplenial cortex (RSGc/RSD), and hippocampus (CA1–3, DG). GiGA1 administration also strongly reduced c-Fos expression in hippocampus and cortex, which has amongst the highest GIRK1 expression^12^ and GiGA1-induced GIRK currents^25^.

We hypothesized that GiGA1 would counteract ethanol-induced hyperactivity in key brain regions implicated in AUD. Indeed, GiGA1 blocked the ethanol-dependent increase in c-Fos expression in the EW, PVT, PVN, and CeA, regions shown previously to exhibit increased neuronal activity in response to ethanol exposure^54^. The EW has been directly linked to alcohol drinking, as genetic manipulation alters consumption, and electrolytic lesions reduce alcohol preference and intake^65,66^. The PVT and PVN regulate hedonic feeding and ethanol consumption^67^. Microinjection studies targeting the PVN show that blocking GABA_A_ receptors reduces voluntary ethanol intake^68^, while stimulation with neuropeptide Y (NPY) enhances ethanol consumption and preference^69^. In the anterior PVT, local injection of orexin enhances ethanol intake, an effect that is blocked by an orexin receptor antagonist^70^. It is possible that action of GiGA1 in the PVN and PVT could contribute to the reduction in sucrose intake observed following GiGA1 administration. Notably, GiGA1 and ethanol produced similar changes in c-Fos expression in regions such as the DLS, RSGc/RSD and hippocampus (CA1–3, DG), while exhibiting opposing effects in the PVT, PVN, and EW, where ethanol typically induces activation and GiGA1 promotes inhibition.

Finally, the CeA is widely recognized as a critical hub in AUD^71^. Lesion studies show that damage to the CeA, but not the neighboring BLA, induces a pronounced decrease in ethanol consumption, while pharmacological studies demonstrate that microinfusions of GABA_A_ receptor antagonists into the CeA suppress drinking behavior^72,73^. The CeA is functionally connected to the NAc, a pathway closely associated with AUD^74,75^. Interestingly, although GiGA1 alone did not significantly affect activity in the CeA or NAc core, it blocked the ethanol-induced increase in c-Fos expression in the CeA and reduced it in the NAc core. However, ethanol did not produce a significant increase in NAc c-Fos expression in our studies, similar to previous work showing that a low dose of ethanol (0.5 g/kg) increases c-Fos expression in the CeA but not in the NAc^54,56^. Additionally, strain differences may play a role, as C57BL/6J mic used in our study show lower c-Fos induction by ethanol in both the core and shell of the NAc compared to DBA/2J mice^57^. The finding that GiGA1 significantly dampened NAc activity alone or in combination with ethanol suggests a broader regulatory effect on the extended amygdala and reward-processing circuits. Together, these findings suggest that GiGA1-dependent GIRK inhibition in these brain regions may play an important role in modulating alcohol-related behaviors.

### Limitations of the study

In the current study, we show that systemic administration of GiGA1 prevents acquisition of ethanol-dependent CPP and reduces ethanol intake in a "drinking in the dark" two-bottle choice self-administration paradigm in both males and females. There are some potential limitations with our study worth considering. First, while systemic application of GiGA1 has clear clinical value, it does not pinpoint where GiGA1 acts in the brain. The brain-wide activity mapping using c-Fos expression, however, did reveal several regions where GiGA1 counteracts the ethanol-activated changes in neuronal activity as inferred from changes in c-Fos levels. Future studies using molecular genetics may reveal more precisely which neurons are affected by GiGA1 in AUD. Second, we focused on GiGA1. However, there are two other published selective activators of GIRK1/GIRK2 channels, ML297 and GAT1508^76,77^. Given the similarity in their pharmacological profiles^25,76,77^, we predict similar results with ML297 and GAT1508. Prior PK studies with GiGA1 indicates that is does not remain elevated for more than 2-3 hours after administration indicating a fast clearance time. Consistent with this, we did not observe any lingering effects of GiGA1 on alcohol drinking (i.e., W6 and W13 EtOH intake were similar even though GiGA1 was administered in between- See **Fig 3B,C**). Lastly, we did not exam c-Fos staining in males. However, the changes in c-Fos expression with ethanol in female mice is remarkably similar to that shown in previously published studies with male mice^54^.

In summary, we show that a selective GIRK1/GIRK2 channel activator has strong therapeutic potential for reducing binge alcohol drinking. Future studies should explore its role in withdrawal management and elucidate the precise neural mechanisms underlying its effects. In addition, assessing GiGA1 in models that more closely capture the chronic features of AUD—such as the chronic intermittent ethanol (CIE) exposure paradigm, which induces escalation of alcohol intake—will be essential to determine its therapeutic potential in more severe contexts.

## Materials and methods

### Mice

All experiments were conducted in accordance with institutional guidelines and an approved protocol by the Institutional Animal Care and Use Committee (IACUC) at the Icahn School of Medicine at Mount Sinai. For all experiments described below, male and female C57BL/6J mice (3–6 months old, approximately 20–35 g; Jackson Laboratory) were used. Mice were housed in groups of 3–5 per cage under a 12-hour light/dark cycle in a temperature- and humidity-controlled environment, with ad libitum access to food and water. Littermates of the same sex were randomly assigned to experimental groups. For 2BC-DID experiments, mice were single-housed two weeks prior to the experiment and placed on a reverse light–dark cycle (lights on at 20:00, lights off at 08:00) to align behavioral testing with their active phase.

### Drug Administration

For the CPP experiments, ethanol (200 proof, ≥99.5%, Sigma-Aldrich, Cat# 493546) was prepared as a 20% v/v solution in 0.9% saline and administered intraperitoneally (IP). The injection volume was adjusted according to body weight, with doses of 1.2 g/kg (7.6 mL/kg) and 2.0 g/kg (12.6 mL/kg). For the 2BC-DID experiments, ethanol was diluted to 10% or 15% (v/v) using the same tap water that mice regularly consumed from their home cage bottles. Ethanol and water were provided in 50 mL Falcon tubes capped with rubber stoppers fitted with drinking tubes (5/16" diameter, Ancare), allowing both bottles to fit within the home cage while replacing the standard single water bottle. GiGA1 was synthesized in collaboration with the National Center for Advancing Translational Sciences (NCATS). The injection solution was prepared by first dissolving GiGA1 (4-8 mg/mL) in dimethyl sulfoxide (DMSO, Sigma-Aldrich, Cat# D8418, final concentration 10%), followed by the addition of polyethylene glycol 300 (PEG300, Laboratory Chemicals, Cat# 0258D, final concentration 40%). Subsequently, Solutol (Kolliphor HS-15, Sigma-Aldrich, Cat# 42996, final concentration 5%) was incorporated at 50°C to maintain solubility and was vortexed to ensure proper mixing. Finally, 0.9% saline (final concentration, 45%) was added, and the mixture was immediately vortexed to ensure that all components were fully dissolved, resulting in a clear and well-incorporated solution. The resulting formulation remained stable for 2–4 hours and was administered intraperitoneally at 5 mL/kg. The vehicle solution was prepared following the same procedure, excluding GiGA1. Baclofen ((±)-Baclofen ≥98% purity by HPLC, Sigma-Aldrich, Cat# B5399) was dissolved in 0.9% saline at 1.5 mg/mL, enabling an IP injection of also 5 mL/kg to achieve a 7.5 mg/kg dose. For Baclofen experiments, control animals were administered 0.9% saline alone as the vehicle.

### Blood Alcohol Concentration (BAC) Measurements

Blood samples were collected via submandibular (facial vein) puncture using sterile Goldenrod lancets for each mouse. Animals were restrained by scruffing at the back of the neck, ensuring that breathing was not compromised, and a quick puncture was made just behind the tip of the mandibular bone. Approximately 50-200 µL of blood was drawn into Microvette 100 Lithium Heparin capillary tubes (Sarstedt), gently tapped to mix with the anticoagulant, and immediately centrifuged at 1000 × g for 10 minutes to separate plasma. The plasma fraction was snap-frozen in liquid nitrogen and stored until batch analysis. All procedures followed recommended guidelines limiting volume to no more than 7.5% of total blood volume on a single occasion for healthy mice. On the day of analysis, plasma ethanol concentration was quantified using an Analox Alcohol Analyzer (GL5 Analyser; Analox Instruments).

### Ethanol-Conditioned-Place Preference (CPP)

For the CPP paradigm, we used a rectangular, custom-built field measuring 64 × 18 cm with a height of 50 cm, divided into three distinct compartments. The two conditioning chambers (26 × 18 cm each) with distinct wall patterns and stainless steel woven wire mesh floors: one featured black and white striped walls and 4.50 mm holes in the floor, whereas the other had gray walls with 1.00 mm holes. The central neutral zone (12 × 18 cm) had a white patterned wall and 1.98 mm holes in the floor. A removable partition allowed each compartment to be isolated during conditioning, and all sessions were conducted during their light phase under red light at 75-125 lux. On Day 1 (pre-test), mice were placed in the center of the apparatus and allowed to freely explore all compartments for 15 minutes to determine their innate side preference. Starting on Day 2, conditioning sessions were performed. Each animal’s least-preferred chamber (identified during the pre-test) was designated as the CS+ and paired with ethanol, whereas the initially favored chamber was paired with saline (CS−) to maximize the detection of ethanol-induced place preference (biased design). Both ethanol and saline were administered immediately before placing the mouse into the corresponding chamber. Two conditioning protocols were used: (Protocol A) a four-day protocol featuring one 5-minute session per day (alternating saline on Days 1 and 3 with ethanol (1.2 g/kg) on Days 2 and 4), and (Protocol B) a five-day protocol in which each day included two 30-minute sessions—a morning saline session and an evening ethanol (2.0 g/kg) session separated by 4–5 hours—for a total of ten sessions. To test GiGA1’s potential to modulate ethanol CPP, we used Protocol A with three treatment groups: (i) vehicle + ethanol, (ii) GiGA1 (30 mg/kg, i.p.) or Baclofen (7.5 mg/kg, i.p.) + saline, and (iii) GiGA1 or Baclofen + ethanol. For ethanol conditioning (groups i and iii), mice received their assigned pre-treatment of vehicle/GiGA1 or Baclofen 30 minutes prior to each ethanol/CS+ session and vehicle 30 minutes before the saline/CS− sessions. For the GiGA1/Baclofen-alone group (ii), mice received pre-treatment 30 minutes before the CS+ sessions, but both CS+ and CS− pairings involved saline instead of ethanol (see Fig. 2B and Fig. 5B). One day after the final conditioning session, a test-day was conducted under the same conditions as the pre-test, during which mice freely explored the apparatus for 15 minutes (see Fig. 1A-B). For cocaine-CPP, cocaine (15 mg/kg) instead of ethanol was used in Protocol A. Time spent in each compartment was recorded, and locomotor activity was measured throughout the protocol. CPP was assessed by statistically comparing the average time on the Paired side with the average time on the Unpaired side on the Test day.

### Novel Object Recognition (NOR) Test

The task was conducted in a square open-field arena under the same environmental conditions as the CPP experiments (light phase, red light at 75–125 lux). The protocol consisted of three consecutive days, with one 10-minute session per day^37^. On Day 1 (habituation), mice were placed individually in the empty arena and allowed to freely explore the field for 10 minutes in the absence of any objects to habituate to the context. On Day 2 (object acquisition), mice received their assigned treatment of vehicle/GiGA1(30 mg/kg, i.p.) 30 minutes prior to the session and were then placed in the arena containing two identical objects positioned symmetrically. Animals were allowed to explore the objects for 10 minutes. Object pairs consisted of a strawberry-shaped toy and a small celery-shaped toy differing in color, shape, and texture, with size (∼one-third of mouse body volume) selected to avoid aversive or anxiety-related responses. To counterbalance object identity and minimize object bias, half of the animals were exposed to one object type during the acquisition session, whereas the remaining animals were exposed to the alternate object type. On Day 3 (test), mice were returned to the arena for a 10-minute session in which one of the familiar objects was replaced by a novel object of the alternate type, such that the familiar and novel objects were counterbalanced across animals. Exploration was defined as active investigation of the object with the nose oriented toward and within close proximity to the object. Locomotor activity was quantified during the acquisition (Day 2) and test (Day 3) sessions, and object exploration time was measured during both sessions. Recognition memory performance was assessed by comparing the time spent exploring the novel versus the familiar object during the test session. A minimum total exploration time (e.g., <5–20 seconds) during testing was used as a filter for analysis.

### Two Bottle Choice – Drinking in the Dark Paradigm (2BC-DID)

All mice underwent a 2-hour daily access to two bottles—one containing water and one containing ethanol—starting 2 hours into the dark cycle, from Monday to Friday. During the first week, the ethanol concentration was 10% (v/v); from the second week onward, it was increased to 15% (v/v). Outside these sessions, mice had continuous access to water. We employed a 2BC-DID paradigm in two Cohorts of mice with subtle differences in their experimental design. In Cohort 1, mice initially conducted their daily 2BC-DID sessions in home cages for six weeks, but beginning in week 7, each mouse was transferred from its home cage to a lickometer cage at the onset of the light cycle (i.e., 14 hours before the 2BC-DID session). Bedding and food from the home cage were transferred to minimize stress, and mice initially had two water bottles to assess side preference. Fifteen minutes before the 2BC-DID session, GiGA1 (30 mg/kg) or vehicle was administered IP. Right before the session, both water bottles were replaced with one fresh water bottle and one ethanol bottle. The ethanol bottle was positioned on the less-preferred side, as determined during the acclimation period, to encourage active ethanol seeking. Lick counts and volume consumed were recorded automatically. Because the lickometer cage setup accommodated one mouse at a time, each animal was tested twice (once with GiGA1 and once with vehicle) on separate days in a counterbalanced order, with the entire Cohort assessed from week 7 to week 12 (see Fig. 3A). Cohort 2 followed the same 2-hour daily access protocol but remained in their home cages throughout. Mice had continuous access to a single water bottle, which was replaced by two bottles (water and ethanol) at the start of each drinking session. Beginning in week 5, vehicle or GiGA1 (30 mg/kg) (or saline/Baclofen, 7.5 mg/kg) was administered immediately before the 2BC-DID session, and total volume consumed from each bottle was measured at the session’s end (see Fig. 4A). Additionally, blood samples were collected in this Cohort to determine blood alcohol concentration (BAC). After the ethanol DID paradigm (water bottle/s replaced by water/4% (w/v) sucrose) in their home cages. In Cohort 1, mice were naïve to 4% (w/v) sucrose prior to testing. In Cohort 2, mice were habituated to 4% (w/v) sucrose for 1 week before testing. In both Cohorts, the treatment was administered at the same time point used for ethanol sessions.

### c-Fos Studies

To mimic the ethanol conditioning process in the CPP paradigm, mice were pretreated with GiGA1 30 minutes before ethanol administration and then placed in a novel open-field environment. This protocol was designed to enhance neuronal activity in regions such as the hippocampus and cortex, which are known to be activated in response to novelty^55,56,78^. We chose to focus on females because of the more robust CPP, higher ethanol BACs, and paucity of studies on females (ref). Female C57BL/6J mice received an IP injection of GiGA1 (30 mg/kg) or vehicle in their home cage, and after 30 minutes, received an IP injection of ethanol (1.2 g/kg) or saline and then placed in a novel open-field environment. After 1 hour, mice were sacrificed, their head removed and immediately placed on ice, while the brains were dissected and then rinsed in cold PBS. Each brain was fixed overnight in 4% paraformaldehyde (PFA) under gentle agitation, then subdivided with a brain matrix into five segments (approximately at Bregma +2, +1, −1, −3 mm along the AP axis), which were post-fixed overnight in 4% PFA to ensure complete penetration. The segments were then embedded in 5% agarose (in PBS) and sectioned into 40-µm slices using a vibratome, which were preserved in PBS containing 0.02% (w/v) sodium azide for further processing. For c-Fos immunostaining, four coronal sections were selected per brain (Bregma: 2.10 mm, 0.98 mm, − 1.22 mm, − 3.08 mm; see Fig. 6A). The regions present in these sections were selected based on their involvement in ethanol-related responses and GIRK1 expression, as reported across studies^49–53^. Free-floating sections were first blocked for 1 hour at room temperature (RT; 100 rpm on an orbital shaker) in PBS (prepared by diluting 10X DPBS, Dulbecco’s formula, Thermo Fisher Scientific, Cat# AAJ61917-K2) containing 0.3% Triton X-100 (Sigma-Aldrich, Cat# X100RS) and 5% normal goat serum (NGS, Thermo Fisher Scientific, Cat# NC0605700). They were then incubated overnight at 4°C with gentle agitation in primary antibody (c-Fos (9F6) Rabbit mAb [1:3000], Cell Signaling Technology Inc, Cat# 2250S) diluted in PBS with 0.3% Triton X-100. After three 5-minute washes in PBS, sections were incubated for 1 hour at RT in the dark with DAPI (0.1 mg/mL final, Sigma-Aldrich, Cat# D9542) and secondary antibody (Invitrogen™ Goat anti-Rabbit IgG (H+L) Highly Cross-Adsorbed Secondary Antibody, Alexa Fluor™ Plus 647 [1:500]; Thermo Fisher Scientific, Cat# PIA32733) diluted in PBS with 0.3% Triton X-100. Sections were then washed again (three 5-minute washes in PBS), mounted onto slides (four sections per slide), air-dried for 15–30 minutes (protected from light), and coverslipped with Fluoromount-G (Thermo Fisher Scientific, Cat# 5018788). Slides were sealed with nail polish and subsequently scanned using a NanoZoomer S60 Digital Slide Scanner (Hamamatsu). A custom macro in FIJI (ImageJ) was used to semi-automatically quantify c-Fos–positive cells in up to 98 brain regions across the four sections.

### Statistical Analyses

All statistical analyses were performed in GraphPad Prism 10. Data are shown as mean ± SEM with individual data points overlaid. Depending on the experimental design, comparisons were made using paired or unpaired t-tests, one-way ANOVA, or two-way ANOVA with appropriate factors (e.g., session, treatment, sex, or their interactions). Significant effects in ANOVA were followed by the indicated multiple comparisons post hoc tests. Exact statistical tests for each figure are reported in the corresponding figure legends. Significance thresholds were set at p < 0.05 (), p < 0.01 (), p < 0.001 (), and p < 0.0001 (****). In Fig. 6, main effects of GiGA1 are additionally denoted with hash symbols (#p < 0.05, ##p < 0.01).

## Supporting information

Taura2026 Supplemental

## Acknowledgements

We thank members of the Slesinger Lab for helpful discussions. We are grateful to Daniel da Silva for assistance with the conditioned place preference (CPP) experiments; the Paul Kenny Lab—Paul Kenny, Stephanie Caligiuri, and Richard O’Connor—for blood alcohol concentration (BAC) measurements; Shen Yao (Endocrinology) for support with the NanoZoomer S60 Digital Slide Scanner; and Katarzyna Cialowicz of the Microscopy and Advanced Bioimaging Core for imaging assistance. This work was supported by NIH grant R01AA018734. The Intramural Research Program of the National Center for Advancing Translational Sciences (NCATS), NIH supported the contributions of SIS and GR to this research under the project # TR00039.

