## Supplementary material for "SELECTIVE ACTIVATION OF GIRK POTASSIUM CHANNELS REDUCES BEHAVIORAL AND BRAIN RESPONSES TO ETHANOL IN MICE": Taura2026 Supplemental

1. Figure S1
2. Figure S2
3. Figure S3
4. Tables S1-S11

**Supplemental Figure Legends:**

**Figure S1. GiGA1 does not alter cocaine-CPP or novel object recognition (relates to Figure 2).**

(A) Cocaine-CPP expressed as time spent in the cocaine-paired side during the Test-day. Mice pretreated with Vehicle or GiGA1 (30 mg/kg, i.p.) 30 min prior to cocaine administration (15 mg/kg; Protocol A) showed robust CPP, with no differences between groups. (B) Novel object recognition (NOR) performance expressed as the discrimination index during the test session. Mice pretreated with Vehicle or GiGA1 (30 mg/kg, i.p.) 30 min prior to the acquisition phase displayed intact recognition memory, with both groups showing greater exploration of the novel object relative to the familiar one. For CPP, main effects of session (pre-test vs. Test-day), treatment (Vehicle vs. GiGA1), and their interaction were analyzed using two-way ANOVA followed by Šídák's multiple comparisons test; paired t-tests compared time spent in saline- vs. cocaine-paired compartments within subjects. For NOR, main effects of object type (familiar vs. novel), treatment, and their interaction were analyzed using two-way ANOVA followed by Holm–Šídák's multiple comparisons test; paired t-tests compared exploration time within subjects. Data are presented as mean ± SEM. *$p < 0.05$, **$p < 0.01$. Sample sizes: CPP, n = 10 per group; NOR, n = 8 per group.

**Figure S2. Dose-response curve for GiGA1 on voluntary ethanol (relates to Figure 2).**

Plot shows ethanol intake plotted as a function of GiGA1 dose (20, 30, 40 mg/kg). N = 15 mice (8 females, 7 males). Data are presented as mean ± SEM. Statistical differences assessed with one-way ANOVA and Šídák's multiple comparisons post hoc test vs vehicle (0 GiGA1). ***$p < 0.0005$; ****$p < 0.0001$.

**Figure S3. Baclofen reduces voluntary ethanol intake also in males (relates to Figure 4).**

(**A–D**) Panels show mean values and individual mice for vehicle and Baclofen sessions for ethanol and water intake (**A,B**), drinking preference (ethanol and water as a proportion of total fluid intake) (**C**), and BAC measurements (**D**). (**E**) Panel shows sucrose (4%) intake for vehicle and Baclofen sessions, after one week of habituation to sucrose consumption under the same drinking regime as used for ethanol in males. N = 16 males were used for the 2BC-DID paradigm. Data are presented as individual mice with mean (bar). Paired t-tests compare vehicle vs. Baclofen treatments. *$p < 0.05$, **$p < 0.01$, ***$p < 0.001$, ****$p < 0.0001$.

**Table S1-S11**: Details of all statistics performed.

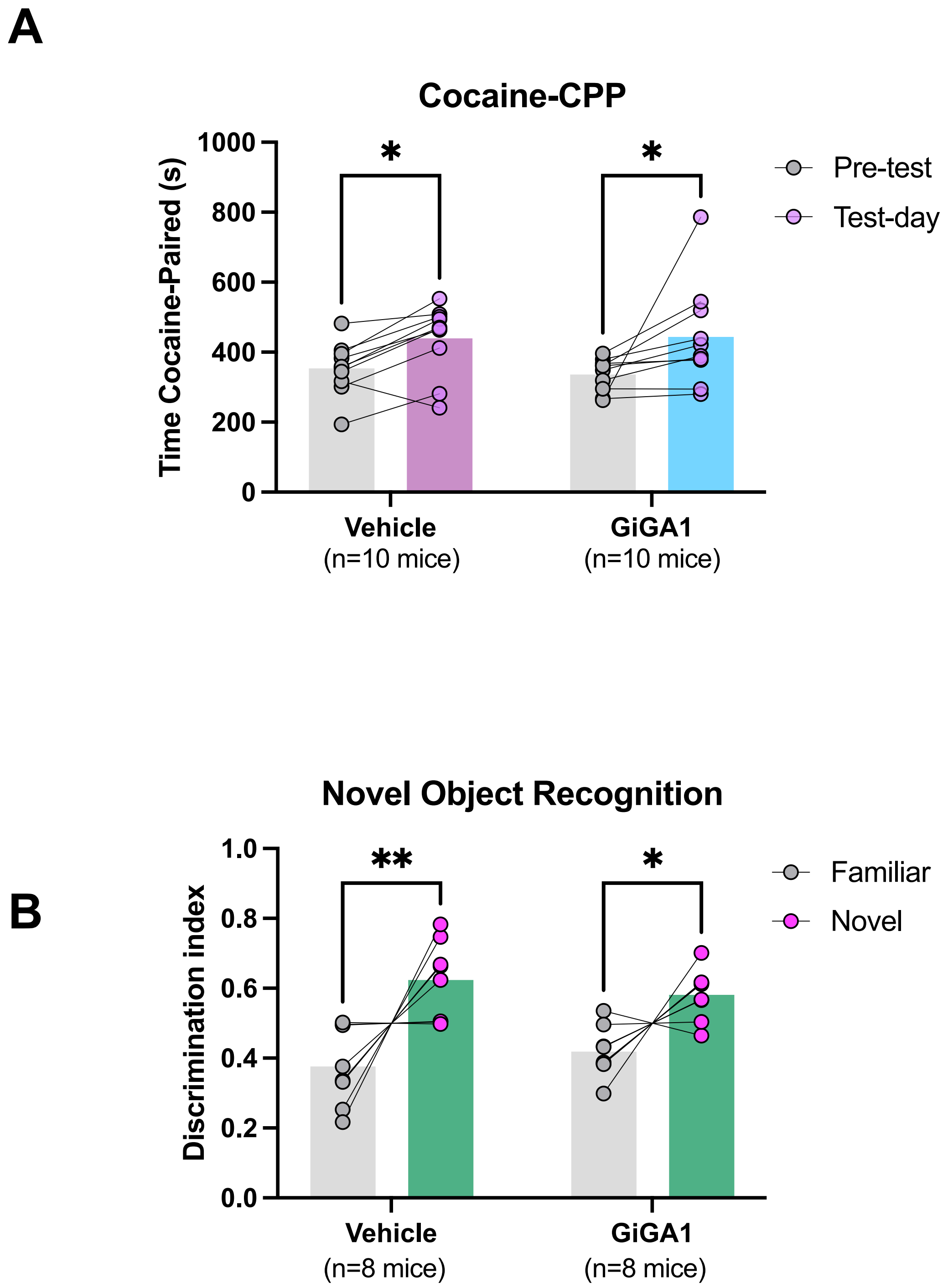
A
Cocaine-CPP
Time Cocaine-Paired (s)
1000
800
600
400
200
0
*
*
Pre-test
Test-day
Vehicle
(n=10 mice)
GiGA1
(n=10 mice)
B
Novel Object Recognition
Discrimination index
1.0
0.8
0.6
0.4
0.2
0.0
**
*
Familiar
Novel
Vehicle
(n=8 mice)
GiGA1
(n=8 mice)

A

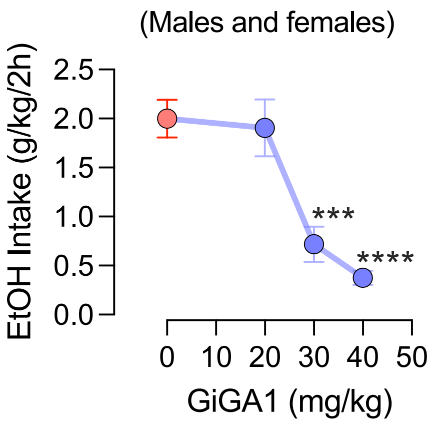

(Males and females)
EtOH Intake (g/kg/2h)
2.5
2.0
1.5
1.0
0.5
0.0
***
****
0
10
20
30
40
50
GiGA1 (mg/kg)

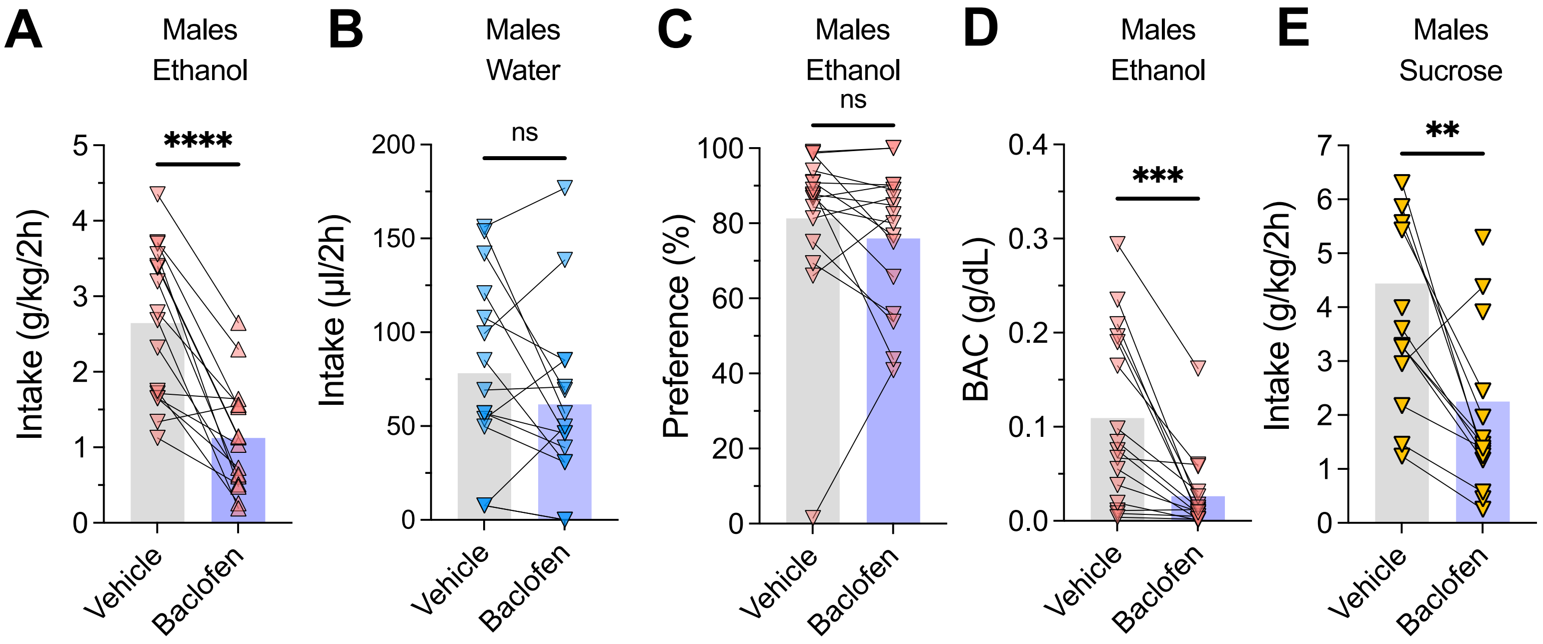
A
Males
Ethanol
****
Intake (g/kg/2h)
0
1
2
3
4
5
Vehicle
Baclofen
B
Males
Water
ns
Intake (µl/2h)
0
50
100
150
200
Vehicle
Baclofen
C
Males
Ethanol
ns
Preference (%)
0
20
40
60
80
100
Vehicle
Baclofen
D
Males
Ethanol
***
BAC (g/dL)
0.0
0.1
0.2
0.3
0.4
Vehicle
Baclofen
E
Males
Sucrose
**
Intake (g/kg/2h)
0
1
2
3
4
5
6
7
Vehicle
Baclofen

**Supplemental Tables S1-S11**

**Table S1 (relates to Figure 1)**

Protocol comparison (Time spent in Paired side)

| **Females** | **N=12** | | | ***Post-hoc Comparison Šídák's*** | | |
|---|---|---|---|---|---|---|
| ***2way-ANOVA*** | *F (DFn, DFd)* | *P value* | | **Pre-test vs Post-test** | *Adjusted P Value* | |
| Protocol x Test | F (1, 22) = 0.8144 | **0.3766** | **ns** | | | |
| Protocol | F (1, 22) = 0.5162 | **0.4800** | **ns** | Protocol A | **0.0122** | * |
| Test | F (1, 22) = 26.92 | **<0.0001** | **####** | Protocol B | **0.0006** | *** |

| **Males** | **N=22** | | | ***Post-hoc Comparison Šídák's*** | | |
|---|---|---|---|---|---|---|
| ***2way-ANOVA*** | *F (DFn, DFd)* | *P value* | | **Pre-test vs Post-test** | *Adjusted P Value* | |
| Protocol x Test | F (1, 42) = 0.7843 | **0.3809** | **ns** | | | |
| Protocol | F (1, 42) = 2.165 | **0.1487** | **ns** | Protocol A | **0.0360** | * |
| Test | F (1, 42) = 19.02 | **0.0001** | **###** | Protocol B | **0.0012** | ** |

Protocol comparison (Distance traveled 0-5min)

| **Females** | **N=12** | | | ***Post-hoc Comparison Šídák's*** | | |
|---|---|---|---|---|---|---|
| ***2way-ANOVA*** | *F (DFn, DFd)* | *P value* | | **Pre-test vs Post-test** | *Adjusted P Value* | |
| Protocol x Test | F (1, 22) = 9.597 | **0.0053** | **##** | | | |
| Ethanol dose | F (1, 22) = 0.02276 | **0.8814** | **#** | Ethanol-1.2g/kg | **0.0178** | * |
| Test | F (1, 22) = 0.9199 | **0.3479** | **ns** | Ethanol-2.0g/kg | **0.2684** | **ns** |

| **Males** | **N=22** | | | ***Post-hoc Comparison Šídák's*** | | |
|---|---|---|---|---|---|---|
| ***2way-ANOVA*** | *F (DFn, DFd)* | *P value* | | **Pre-test vs Post-test** | *Adjusted P Value* | |
| Protocol x Test | F (1, 42) = 5.220 | **0.0275** | **#** | | | |
| Ethanol dose | F (1, 42) = 10.72 | **0.0021** | **##** | Ethanol-1.2g/kg | **0.9882** | **ns** |
| Test | F (1, 42) = 6.144 | **0.0173** | **#** | Ethanol-2.0g/kg | **0.0033** | ** |

Protocol comparison (Distance traveled 25-30min)

| ***Paired t-test*** | *Group* | *N* | *t, df* | *P value* | |
|---|---|---|---|---|---|
| Ethanol-2.0g/kg vs Saline | Females | 12 | t=6.528, df=11 | **<0.0001** | **** |

| ***Paired t-test*** | *Group* | *N* | *t, df* | *P value* | |
|---|---|---|---|---|---|
| Ethanol-2.0g/kg vs Saline | Males | 22 | t=5.765, df=21 | **<0.0001** | **** |

**Table S2 (relates to Figure 2)**

GiGA1 treatment (Time spent in Paired side)

| **Females** | **N=12** | | | *Post-hoc Comparison Šídák's* | | |
|---|---|---|---|---|---|---|
| ***2way-ANOVA*** | *F (DFn, DFd)* | *P value* | | **Pre-test vs Post-test** | *Adjusted P Value* | |
| Treatment x Test | F (2, 33) = 3.837 | **0.0318** | **#** | Vehicle + Ethanol | **0.0157** | ***** |
| Treatment | F (2, 33) = 3.395 | **0.0456** | **#** | GiGA1 + Saline | **0.7866** | **ns** |
| Test | F (1, 33) = 5.068 | **0.0311** | **#** | GiGA1 + Ethanol | **0.2405** | **ns** |

| **Males** | **N=10-26** | | | *Post-hoc Comparison Šídák's* | | |
|---|---|---|---|---|---|---|
| ***2way-ANOVA*** | *F (DFn, DFd)* | *P value* | | **Pre-test vs Post-test** | *Adjusted P Value* | |
| Treatment x Test | F (2, 57) = 4.078 | **0.0221** | **#** | Vehicle + Ethanol | **0.0133** | ***** |
| Treatment | F (2, 57) = 5.332 | **0.0075** | **##** | GiGA1 + Saline | **0.4208** | **ns** |
| Test | F (1, 57) = 2.240 | **0.1400** | **ns** | GiGA1 + Ethanol | **0.0527** | **ns** |

GiGA1 treatment (Distance traveled)

| **Females** | **N=12** | | | *Post-hoc Comparison Šídák's* | | |
|---|---|---|---|---|---|---|
| ***2way-ANOVA*** | *F (DFn, DFd)* | *P value* | | **Unpaired vs Paired** | *Adjusted P Value* | |
| Treatment x Test | F (2, 33) = 19.49 | **<0.0001** | **####** | Vehicle + Ethanol | **0.0238** | ***** |
| Treatment | F (1, 33) = 15.09 | **0.0005** | **###** | GiGA1 + Saline | **<0.0001** | ******** |
| Test | F (2, 33) = 8.256 | **0.0012** | **##** | GiGA1 + Ethanol | **0.0005** | ******* |

| **Males** | **N=10-26** | | | *Post-hoc Comparison Šídák's* | | |
|---|---|---|---|---|---|---|
| ***2way-ANOVA*** | *F (DFn, DFd)* | *P value* | | **Unpaired vs Paired** | *Adjusted P Value* | |
| Treatment x Test | F (2, 57) = 7.430 | **0.0014** | **##** | Vehicle + Ethanol | **0.9492** | **ns** |
| Treatment | F (1, 57) = 30.12 | **<0.0001** | **####** | GiGA1 + Saline | **0.0015** | ****** |
| Test | F (2, 57) = 4.472 | **0.0157** | **#** | GiGA1 + Ethanol | **<0.0001** | ******** |

**Table S3 (relates to Figure 3)**

Intake

| ***Paired t-test*** | *Group* | *N* | *t, df* | *P value* | |
|---|---|---|---|---|---|
| GiGA1-30mg/kg vs Vehicle | Ethanol | 15 | t=3.810, df=14 | **0.0019** | ****** |

| ***Paired t-test*** | *Group* | *N* | *t, df* | *P value* | |
|---|---|---|---|---|---|
| GiGA1-30mg/kg vs Vehicle | Water | 15 | t=0.3078, df=14 | **0.7628** | **ns** |

Licks

| *Paired t-test* | *Group* | *N* | *t, df* | *P value* | |
|---|---|---|---|---|---|
| GiGA1-30mg/kg vs Vehicle | Ethanol | 15 | t=3.233, df=14 | **0.0060** | ** |

| *Paired t-test* | *Group* | *N* | *t, df* | *P value* | |
|---|---|---|---|---|---|
| GiGA1-30mg/kg vs Vehicle | Water | 15 | t=1.427, df=14 | **0.1754** | **ns** |

Preference

| *Paired t-test* | *Group* | *N* | *t, df* | *P value* | |
|---|---|---|---|---|---|
| GiGA1-30mg/kg vs Vehicle | Ethanol | 15 | t=1.676, df=14 | **0.1159** | **ns** |

Sucrose intake

| *Paired t-test* | *Group* | *N* | *t, df* | *P value* | |
|---|---|---|---|---|---|
| GiGA1-30mg/kg vs Vehicle | Sucrose | 12 | t=0.4957, df=11 | **0.6299** | **ns** |

**Table S4 (relates to Figure 4)**

Ethanol intake and preference time course

| **Ethanol intake** | **N=16** | | | ***Post-hoc Comparison*** | ***Šídák's*** | |
|---|---|---|---|---|---|---|
| ***2way-ANOVA*** | *F (DFn, DFd)* | *P value* | | **Females vs Males** | *Adjusted P Value* | |
| Time x Sex | F (3, 90) = 3.002 | **0.0346** | **##** | Week 1 | **0.5819** | **ns** |
| Time | F (3, 90) = 67.16 | **<0.0001** | **####** | Week 2 | **0.0165** | * |
| Sex | F (1, 30) = 9.420 | **0.0045** | **##** | Week 3 | **0.0454** | * |
| | | | | Week 4 | **0.0010** | ** |

| **Preference** | **N=16** | | | ***Post-hoc Comparison*** | ***Šídák's*** | |
|---|---|---|---|---|---|---|
| ***2way-ANOVA*** | *F (DFn, DFd)* | *P value* | | **Females vs Males** | *Adjusted P Value* | |
| Time x Sex | F (3, 90) = 0.3487 | **0.7901** | **ns** | Week 1 | **0.0980** | **ns** |
| Time | F (3, 90) = 12.58 | **<0.0001** | **####** | Week 2 | **0.6678** | **ns** |
| Sex | F (1, 30) = 5.394 | **0.0272** | **#** | Week 3 | **0.6713** | **ns** |
| | | | | Week 4 | **0.4321** | **ns** |

Intake GiGA1 treatment (Females)

| *Paired t-test* | *Group* | *N* | *t, df* | *P value* | |
|---|---|---|---|---|---|
| GiGA1-30mg/kg vs Vehicle | Ethanol | 16 | t=3.985, df=15 | **0.0012** | ** |

| *Paired t-test* | *Group* | *N* | *t, df* | *P value* | |
|---|---|---|---|---|---|
| GiGA1-30mg/kg vs Vehicle | Water | 16 | t=0.3156, df=15 | **0.7566** | **ns** |

Intake GiGA1 treatment (Males)

| | ***Paired t-test*** | *Group* | *N* | *t, df* | *P value* | |
|---|---|---|---|---|---|---|
| | GiGA1-30mg/kg vs Vehicle | Ethanol | 16 | t=3.190, df=15 | **0.0061** | ** |

| | ***Paired t-test*** | *Group* | *N* | *t, df* | *P value* | |
|---|---|---|---|---|---|---|
| | GiGA1-30mg/kg vs Vehicle | Water | 16 | t=0.7035, df=15 | **0.4925** | **ns** |

Preference GiGA1 treatment

| | ***Paired t-test*** | *Group* | *N* | *t, df* | *P value* | |
|---|---|---|---|---|---|---|
| | GiGA1-30mg/kg vs Vehicle | Females | 16 | t=3.551, df=15 | **0.0029** | ** |

| | ***Paired t-test*** | *Group* | *N* | *t, df* | *P value* | |
|---|---|---|---|---|---|---|
| | GiGA1-30mg/kg vs Vehicle | Males | 16 | t=1.736, df=15 | **0.1030** | **ns** |

BAC GiGA1 treatment

| | ***Paired t-test*** | *Group* | *N* | *t, df* | *P value* | |
|---|---|---|---|---|---|---|
| | GiGA1-30mg/kg vs Vehicle | Females | 16 | t=4.067, df=15 | **0.0010** | ** |

| | ***Paired t-test*** | *Group* | *N* | *t, df* | *P value* | |
|---|---|---|---|---|---|---|
| | GiGA1-30mg/kg vs Vehicle | Males | 14 | t=3.431, df=13 | **0.0045** | ** |

Sucrose intake GiGA1 treatment

| | ***Paired t-test*** | *Group* | *N* | *t, df* | *P value* | |
|---|---|---|---|---|---|---|
| | GiGA1-30mg/kg vs Vehicle | Females | 16 | t=2.948, df=15 | **0.0100** | ** |

| | ***Paired t-test*** | *Group* | *N* | *t, df* | *P value* | |
|---|---|---|---|---|---|---|
| | GiGA1-30mg/kg vs Vehicle | Males | 16 | t=3.783, df=15 | **0.0018** | ** |

**Table S5 (relates to Figure 5)**

Baclofen treatment (Time spent in Paired side)

| **Females** | **N=12** | | | ***Post-hoc Comparison Šídák's*** | | |
|---|---|---|---|---|---|---|
| ***2way-ANOVA*** | *F (DFn, DFd)* | *P value* | | **Unpaired vs Paired** | *Adjusted P Value* | |
| Treatment x Test | F (2, 44) = 1.378 | **0.2628** | **ns** | Vehicle + Ethanol | **0.0063** | ** |
| Treatment | F (2, 44) = 0.4232 | **0.6576** | **ns** | Baclofen + Saline | **0.5681** | **ns** |
| Test | F (1, 44) = 18.95 | **<0.0001** | **####** | Baclofen + Ethanol | **0.0109** | * |

Baclofen treatment (Distance traveled)

| **Females** | **N=12** | | | ***Post-hoc Comparison Šídák's*** | | |
|---|---|---|---|---|---|---|
| ***2way-ANOVA*** | *F (DFn, DFd)* | *P value* | | **Unpaired vs Paired** | *Adjusted P Value* | |
| Treatment x Test | F (2, 44) = 21.65 | <0.0001 | #### | Vehicle + Ethanol | **0.0249** | * |
| Treatment | F (1, 44) = 7.121 | 0.0106 | ## | Baclofen + Saline | **<0.0001** | **** |
| Test | F (2, 44) = 3.616 | 0.0352 | # | Baclofen + Ethanol | **0.7425** | **ns** |

Intake Baclofen treatment (Females)

| ***Paired t-test*** | *Group* | *N* | *t, df* | *P value* | |
|---|---|---|---|---|---|
| Baclofen-7.5mg/kg vs Vehicle | Ethanol | 15 | t=10.48, df=14 | **<0.0001** | **** |

| ***Paired t-test*** | *Group* | *N* | *t, df* | *P value* | |
|---|---|---|---|---|---|
| Baclofen-7.5mg/kg vs Vehicle | Water | 16 | t=0.1297, df=15 | **0.8985** | **ns** |

Preference Baclofen treatment (Females)

| ***Paired t-test*** | *Group* | *N* | *t, df* | *P value* | |
|---|---|---|---|---|---|
| Baclofen-7.5mg/kg vs Vehicle | Ethanol | 16 | t=1.124, df=15 | **0.2788** | **ns** |

BAC Baclofen treatment

| ***Paired t-test*** | *Group* | *N* | *t, df* | *P value* | |
|---|---|---|---|---|---|
| Baclofen-7.5mg/kg vs Vehicle | Females | 16 | t=4.681, df=15 | **0.0003** | *** |

Sucrose intake Baclofen treatment

| ***Paired t-test*** | *Group* | *N* | *t, df* | *P value* | |
|---|---|---|---|---|---|
| Baclofen-7.5mg/kg vs Vehicle | Females | 16 | t=3.691, df=15 | **0.0022** | ** |

**Table S6 (relates to Figure 7)**

c-Fos activity

| **Pir(r)** | **N=18** | | | ***Post-hoc Comparison Šídák's*** | | |
|---|---|---|---|---|---|---|
| ***2way-ANOVA*** | *F (DFn, DFd)* | *P value* | | | *Adjusted P Value* | |
| Interaction | F (1, 68) = 7.127 | **0.0095** | **##** | Saline+Vehicle vs Saline+GiGA1 | **0.4583** | **ns** |
| Ethanol-treatment | F (1, 68) = 3.382 | **0.0703** | **ns** | Saline+Vehicle vs Ethanol+Vehicle | **0.0065** | ** |
| GiGA1-treatment | F (1, 68) = 20.83 | **<0.0001** | **####** | Ethanol+Vehicle vs Ethanol+GiGA1 | **<0.0001** | **** |

| VTT | N=18 | | | Post-hoc Comparison Šidák's | | |
|---|---|---|---|---|---|---|
| *2way-ANOVA* | *F (DFn, DFd)* | *P value* | | | *Adjusted P Value* | |
| Interaction | F (1, 68) = 1.754 | **0.1899** | **ns** | Saline+Vehicle vs Saline+GiGA1 | **0.5275** | **ns** |
| Ethanol-treatment | F (1, 68) = 1.013 | **0.3177** | **ns** | Saline+Vehicle vs Ethanol+Vehicle | **0.2805** | **ns** |
| GiGA1-treatment | F (1, 68) = 9.429 | **0.0031** | **##** | Ethanol+Vehicle vs Ethanol+GiGA1 | **0.0082** | ** |

| RSGc | N=18 | | | Post-hoc Comparison Šidák's | | |
|---|---|---|---|---|---|---|
| *2way-ANOVA* | *F (DFn, DFd)* | *P value* | | | *Adjusted P Value* | |
| Interaction | F (1, 68) = 3.383 | **0.0703** | **ns** | Saline+Vehicle vs Saline+GiGA1 | **0.0093** | ** |
| Ethanol-treatment | F (1, 68) = 4.797 | **0.0319** | **#** | Saline+Vehicle vs Ethanol+Vehicle | **0.0173** | * |
| GiGA1-treatment | F (1, 68) = 6.230 | **0.0150** | **#** | Ethanol+Vehicle vs Ethanol+GiGA1 | **0.9548** | **ns** |

| BMA | N=18 | | | Post-hoc Comparison Šidák's | | |
|---|---|---|---|---|---|---|
| *2way-ANOVA* | *F (DFn, DFd)* | *P value* | | | *Adjusted P Value* | |
| Interaction | F (1, 68) = 0.05881 | **0.8091** | **ns** | Saline+Vehicle vs Saline+GiGA1 | **0.6873** | **ns** |
| Ethanol-treatment | F (1, 68) = 0.8180 | **0.3690** | **ns** | Saline+Vehicle vs Ethanol+Vehicle | **0.9538** | **ns** |
| GiGA1-treatment | F (1, 68) = 2.741 | **0.1024** | **ns** | Ethanol+Vehicle vs Ethanol+GiGA1 | **0.4567** | **ns** |

| CeA | N=18 | | | Post-hoc Comparison Šidák's | | |
|---|---|---|---|---|---|---|
| *2way-ANOVA* | *F (DFn, DFd)* | *P value* | | | *Adjusted P Value* | |
| Interaction | F (1, 68) = 3.098 | **0.0829** | **ns** | Saline+Vehicle vs Saline+GiGA1 | **0.3165** | **ns** |
| Ethanol-treatment | F (1, 68) = 9.680 | **0.0027** | **##** | Saline+Vehicle vs Ethanol+Vehicle | **0.0029** | ** |
| GiGA1-treatment | F (1, 68) = 0.2229 | **0.6383** | **ns** | Ethanol+Vehicle vs Ethanol+GiGA1 | **0.7448** | **ns** |

| DG | N=18 | | | Post-hoc Comparison Šidák's | | |
|---|---|---|---|---|---|---|
| *2way-ANOVA* | *F (DFn, DFd)* | *P value* | | | *Adjusted P Value* | |
| Interaction | F (1, 68) = 4.476 | **0.0380** | **#** | Saline+Vehicle vs Saline+GiGA1 | **0.0270** | * |
| Ethanol-treatment | F (1, 68) = 2.977 | **0.0890** | **ns** | Saline+Vehicle vs Ethanol+Vehicle | **0.0249** | * |

| | | | | | | |
|---|---|---|---|---|---|---|
| GiGA1-treatment | F (1, 68) = 2.831 | **0.0971** | **ns** | Ethanol+Vehicle vs Ethanol+GiGA1 | **0.9862** | **ns** |

| **CA3** | **N=18** | | | ***Post-hoc Comparison Šídák's*** | | |
|---|---|---|---|---|---|---|
| ***2way-ANOVA*** | *F (DFn, DFd)* | *P value* | | | *Adjusted P Value* | |
| Interaction | F (1, 68) = 3.021 | **0.0867** | **ns** | Saline+Vehicle vs Saline+GiGA1 | **<0.0001** | ******** |
| Ethanol-treatment | F (1, 68) = 13.68 | **0.0004** | **###** | Saline+Vehicle vs Ethanol+Vehicle | **0.0008** | ******* |
| GiGA1-treatment | F (1, 68) = 20.75 | **<0.0001** | **####** | Ethanol+Vehicle vs Ethanol+GiGA1 | **0.1437** | **ns** |

| **DLS** | **N=18** | | | ***Post-hoc Comparison Šídák's*** | | |
|---|---|---|---|---|---|---|
| ***2way-ANOVA*** | *F (DFn, DFd)* | *P value* | | | *Adjusted P Value* | |
| Interaction | F (1, 68) = 3.498 | **0.0657** | **ns** | Saline+Vehicle vs Saline+GiGA1 | **0.0011** | ****** |
| Ethanol-treatment | F (1, 68) = 13.13 | **0.0006** | **###** | Saline+Vehicle vs Ethanol+Vehicle | **0.0007** | ******* |
| GiGA1-treatment | F (1, 68) = 11.83 | **0.0010** | **##** | Ethanol+Vehicle vs Ethanol+GiGA1 | **0.6126** | **ns** |

| **EW** | **N=18** | | | ***Post-hoc Comparison Šídák's*** | | |
|---|---|---|---|---|---|---|
| ***2way-ANOVA*** | *F (DFn, DFd)* | *P value* | | | *Adjusted P Value* | |
| Interaction | F (1, 68) = 3.412 | **0.0691** | **ns** | Saline+Vehicle vs Saline+GiGA1 | **0.2529** | **ns** |
| Ethanol-treatment | F (1, 68) = 16.46 | **0.0001** | **###** | Saline+Vehicle vs Ethanol+Vehicle | **0.0003** | ******* |
| GiGA1-treatment | F (1, 68) = 18.14 | **<0.0001** | **####** | Ethanol+Vehicle vs Ethanol+GiGA1 | **0.0002** | ******* |

| **NAcC** | **N=18** | | | ***Post-hoc Comparison Šídák's*** | | |
|---|---|---|---|---|---|---|
| ***2way-ANOVA*** | *F (DFn, DFd)* | *P value* | | | *Adjusted P Value* | |
| Interaction | F (1, 68) = 2.102 | **0.1517** | **ns** | Saline+Vehicle vs Saline+GiGA1 | **0.6053** | **ns** |
| Ethanol-treatment | F (1, 68) = 0.4987 | **0.4825** | **ns** | Saline+Vehicle vs Ethanol+Vehicle | **0.3460** | **ns** |
| GiGA1-treatment | F (1, 68) = 9.207 | **0.0034** | **##** | Ethanol+Vehicle vs Ethanol+GiGA1 | **0.0068** | ****** |

| **PVT** | **N=18** | | | ***Post-hoc Comparison Šídák's*** | | |
|---|---|---|---|---|---|---|
| ***2way-ANOVA*** | *F (DFn, DFd)* | *P value* | | | *Adjusted P Value* | |
| Interaction | F (1, 68) = 8.350 | **0.0052** | **##** | Saline+Vehicle vs Saline+GiGA1 | **0.9995** | **ns** |

| | | | | | | |
|---|---|---|---|---|---|---|
| Ethanol-treatment | F (1, 68) = 12.28 | **0.0008** | **###** | Saline+Vehicle vs Ethanol+Vehicle | **<0.0001** | ******** |
| GiGA1-treatment | F (1, 68) = 9.178 | **0.0035** | **##** | Ethanol+Vehicle vs Ethanol+GiGA1 | **0.0003** | ******* |

| **PVN** | **N=18** | | | ***Post-hoc Comparison Šídák's*** | | |
|---|---|---|---|---|---|---|
| ***2way-ANOVA*** | *F (DFn, DFd)* | *P value* | | | *Adjusted P Value* | |
| Interaction | F (1, 68) = 3.982 | **0.0500** | **#** | Saline+Vehicle vs Saline+GiGA1 | **0.6800** | **ns** |
| Ethanol-treatment | F (1, 68) = 3.565 | **0.0633** | **ns** | Saline+Vehicle vs Ethanol+Vehicle | **0.0230** | ***** |
| GiGA1-treatment | F (1, 68) = 11.72 | **0.0010** | **##** | Ethanol+Vehicle vs Ethanol+GiGA1 | **0.0008** | ******* |

**Table S7 (relates to Figure 7M)**

c-Fos activity heatmap

| **N=18** | ***Post-hoc Comparison*** | ***Šídák's*** | **Summary** |
|---|---|---|---|
| ***2way-ANOVA*** | *Fold-change* | *Adjusted P Value* | |
| **Pir(r)** | | | |
| Saline vs. Ethanol-1.2g/kg | 0.4577 | **0.0459** | ● |
| Saline vs. GiGA1-30mg/Kg | -0.1924 | **0.6700** | **ns** |
| Saline vs. GiGA1+Ethanol | -0.2769 | **0.3709** | **ns** |
| **Pir(rc)** | | | |
| Saline vs. Ethanol-1.2g/kg | -0.0971 | **0.9395** | **ns** |
| Saline vs. GiGA1-30mg/Kg | -0.1149 | **0.9047** | **ns** |
| Saline vs. GiGA1+Ethanol | -0.1303 | **0.8680** | **ns** |
| **Pir(c)** | | | |
| Saline vs. Ethanol-1.2g/kg | -0.1661 | **0.7615** | **ns** |
| Saline vs. GiGA1-30mg/Kg | -0.1528 | **0.8038** | **ns** |
| Saline vs. GiGA1+Ethanol | -0.3657 | **0.1512** | **ns** |
| **VTT** | | | |
| Saline vs. Ethanol-1.2g/kg | 0.2879 | **0.3368** | **ns** |
| Saline vs. GiGA1-30mg/Kg | -0.2159 | **0.5842** | **ns** |
| Saline vs. GiGA1+Ethanol | -0.2553 | **0.4425** | **ns** |
| **PrL** | | | |
| Saline vs. Ethanol-1.2g/kg | -0.0796 | **0.9653** | **ns** |
| Saline vs. GiGA1-30mg/Kg | -0.0671 | **0.9787** | **ns** |

| | | | |
|---|---|---|---|
| Saline vs. GiGA1+Ethanol | -0.0662 | **0.9795** | **ns** |
| **Ect(r)** | | | |
| Saline vs. Ethanol-1.2g/kg | -0.3082 | **0.2786** | **ns** |
| Saline vs. GiGA1-30mg/Kg | -0.2416 | **0.4906** | **ns** |
| Saline vs. GiGA1+Ethanol | -0.4507 | **0.0507** | **ns** |
| | 0.0000 | | |
| **Ect(c)** | 0.0000 | | |
| Saline vs. Ethanol-1.2g/kg | 0.0101 | **>0.9999** | **ns** |
| Saline vs. GiGA1-30mg/Kg | -0.2462 | **0.4743** | **ns** |
| Saline vs. GiGA1+Ethanol | -0.0862 | **0.9565** | **ns** |
| **M2** | | | |
| Saline vs. Ethanol-1.2g/kg | -0.1622 | **0.7741** | **ns** |
| Saline vs. GiGA1-30mg/Kg | -0.3617 | **0.1583** | **ns** |
| Saline vs. GiGA1+Ethanol | -0.5786 | **0.0067** | ●● |
| **Cg(r)** | | | |
| Saline vs. Ethanol-1.2g/kg | -0.1141 | **0.9065** | **ns** |
| Saline vs. GiGA1-30mg/Kg | -0.0206 | **0.9994** | **ns** |
| Saline vs. GiGA1+Ethanol | 0.0101 | **>0.9999** | **ns** |
| **Cg c** | | | |
| Saline vs. Ethanol-1.2g/kg | -0.2279 | **0.5402** | **ns** |
| Saline vs. GiGA1-30mg/Kg | -0.1824 | **0.7056** | **ns** |
| Saline vs. GiGA1+Ethanol | -0.2934 | **0.3204** | **ns** |
| **RSGc** | | | |
| Saline vs. Ethanol-1.2g/kg | -0.4959 | **0.0261** | ● |
| Saline vs. GiGA1-30mg/Kg | -0.5334 | **0.0144** | ● |
| Saline vs. GiGA1+Ethanol | -0.5767 | **0.0069** | ●● |
| **RSD** | | | |
| Saline vs. Ethanol-1.2g/kg | -0.4438 | **0.0559** | **ns** |
| Saline vs. GiGA1-30mg/Kg | -0.4206 | **0.0765** | **ns** |
| Saline vs. GiGA1+Ethanol | -0.5238 | **0.0168** | ● |
| **BLA** | | | |
| Saline vs. Ethanol-1.2g/kg | -0.0860 | **0.9568** | **ns** |
| Saline vs. GiGA1-30mg/Kg | -0.1739 | **0.7349** | **ns** |
| Saline vs. GiGA1+Ethanol | -0.1627 | **0.7724** | **ns** |

| | | | |
|---|---|---|---|
| **BMA** | | | |
| Saline vs. Ethanol-1.2g/kg | -0.0856 | **0.9574** | **ns** |
| Saline vs. GiGA1-30mg/Kg | -0.1829 | **0.7037** | **ns** |
| Saline vs. GiGA1+Ethanol | -0.3312 | **0.2211** | **ns** |
| | 0.0000 | | |
| **LaL** | 0.0000 | | |
| Saline vs. Ethanol-1.2g/kg | -0.0498 | **0.9910** | **ns** |
| Saline vs. GiGA1-30mg/Kg | 0.0046 | **>0.9999** | **ns** |
| Saline vs. GiGA1+Ethanol | 0.0041 | **>0.9999** | **ns** |
| **CeA** | | | |
| Saline vs. Ethanol-1.2g/kg | 0.7628 | **0.0002** | ●●● |
| Saline vs. GiGA1-30mg/Kg | 0.3495 | **0.1816** | **ns** |
| Saline vs. GiGA1+Ethanol | 0.5611 | **0.0091** | ●● |
| **MeP** | | | |
| Saline vs. Ethanol-1.2g/kg | 0.1343 | **0.8574** | **ns** |
| Saline vs. GiGA1-30mg/Kg | -0.0019 | **>0.9999** | **ns** |
| Saline vs. GiGA1+Ethanol | -0.0611 | **0.9838** | **ns** |
| **DG** | | | |
| Saline vs. Ethanol-1.2g/kg | -0.2930 | **0.3216** | **ns** |
| Saline vs. GiGA1-30mg/Kg | -0.2898 | **0.3311** | **ns** |
| Saline vs. GiGA1+Ethanol | -0.2599 | **0.4267** | **ns** |
| **CA12** | | | |
| Saline vs. Ethanol-1.2g/kg | -0.6391 | **0.0022** | ●● |
| Saline vs. GiGA1-30mg/Kg | -0.5458 | **0.0117** | ● |
| Saline vs. GiGA1+Ethanol | -0.8030 | **<0.0001** | ●●●● |
| **CA3** | | | |
| Saline vs. Ethanol-1.2g/kg | -0.3998 | **0.1001** | **ns** |
| Saline vs. GiGA1-30mg/Kg | -0.4628 | **0.0427** | ● |
| Saline vs. GiGA1+Ethanol | -0.6073 | **0.0040** | ●● |
| **DMS** | | | |
| Saline vs. Ethanol-1.2g/kg | -0.2862 | **0.3419** | **ns** |
| Saline vs. GiGA1-30mg/Kg | -0.2743 | **0.3791** | **ns** |
| Saline vs. GiGA1+Ethanol | -0.5028 | **0.0234** | ● |
| **DLS** | | | |
| Saline vs. Ethanol-1.2g/kg | -0.5736 | **0.0073** | ●● |

| | | | |
|---|---|---|---|
| Saline vs. GiGA1-30mg/Kg | -0.5544 | **0.0101** | ● |
| Saline vs. GiGA1+Ethanol | -0.7376 | **0.0003** | ●●● |
| **VMS** | | | |
| Saline vs. Ethanol-1.2g/kg | -0.2631 | **0.4160** | **ns** |
| Saline vs. GiGA1-30mg/Kg | -0.2561 | **0.4398** | **ns** |
| Saline vs. GiGA1+Ethanol | -0.1812 | **0.7099** | **ns** |
| **VLS** | | | |
| Saline vs. Ethanol-1.2g/kg | -0.2655 | **0.4080** | **ns** |
| Saline vs. GiGA1-30mg/Kg | -0.4762 | **0.0351** | ● |
| Saline vs. GiGA1+Ethanol | 0.8246 | **<0.0001** | ●●●● |
| **NAcC** | | | |
| Saline vs. Ethanol-1.2g/kg | 0.2904 | **0.3291** | **ns** |
| Saline vs. GiGA1-30mg/Kg | -0.2134 | **0.5931** | **ns** |
| Saline vs. GiGA1+Ethanol | -0.3137 | **0.2641** | **ns** |
| **LAcbSh** | | | |
| Saline vs. Ethanol-1.2g/kg | -0.1365 | **0.8514** | **ns** |
| Saline vs. GiGA1-30mg/Kg | -0.3062 | **0.2842** | **ns** |
| Saline vs. GiGA1+Ethanol | -0.3530 | **0.1747** | **ns** |
| **MAcbSh** | | | |
| Saline vs. Ethanol-1.2g/kg | 0.2153 | **0.5864** | **ns** |
| Saline vs. GiGA1-30mg/Kg | -0.2018 | **0.6357** | **ns** |
| Saline vs. GiGA1+Ethanol | -0.2066 | **0.6182** | **ns** |
| **PVT** | | | |
| Saline vs. Ethanol-1.2g/kg | 0.7923 | **<0.0001** | ●●●● |
| Saline vs. GiGA1-30mg/Kg | -0.0173 | **0.9996** | **ns** |
| Saline vs. GiGA1+Ethanol | 0.0588 | **0.9855** | **ns** |
| **LDVL** | | | |
| Saline vs. Ethanol-1.2g/kg | -0.3964 | **0.1044** | **ns** |
| Saline vs. GiGA1-30mg/Kg | -0.4850 | **0.0308** | ● |
| Saline vs. GiGA1+Ethanol | -0.6822 | **0.0009** | ●●● |
| **VL** | | | |
| Saline vs. Ethanol-1.2g/kg | -0.4290 | **0.0684** | **ns** |
| Saline vs. GiGA1-30mg/Kg | -0.4169 | **0.0804** | **ns** |
| Saline vs. GiGA1+Ethanol | -0.6664 | **0.0013** | ●● |

| | | | |
|---|---|---|---|
| **PVN** | | | |
| Saline vs. Ethanol-1.2g/kg | 0.4326 | **0.0651** | **ns** |
| Saline vs. GiGA1-30mg/Kg | -0.1591 | **0.7843** | **ns** |
| Saline vs. GiGA1+Ethanol | -0.1711 | **0.7448** | **ns** |

| | | | |
|---|---|---|---|
| PLH | | | |
| Saline vs. Ethanol-1.2g/kg | -0.0972 | **0.9395** | **ns** |
| Saline vs. GiGA1-30mg/Kg | -0.0368 | **0.9963** | **ns** |
| Saline vs. GiGA1+Ethanol | -0.1267 | **0.8770** | **ns** |

| | | | |
|---|---|---|---|
| **VMHC** | | | |
| Saline vs. Ethanol-1.2g/kg | -0.1590 | **0.7844** | **ns** |
| Saline vs. GiGA1-30mg/Kg | 0.0039 | **>0.9999** | **ns** |
| Saline vs. GiGA1+Ethanol | -0.0832 | **0.9607** | **ns** |

| | | | |
|---|---|---|---|
| **VTA** | | | |
| Saline vs. Ethanol-1.2g/kg | -0.2023 | **0.6339** | **ns** |
| Saline vs. GiGA1-30mg/Kg | -0.0881 | **0.9538** | **ns** |
| Saline vs. GiGA1+Ethanol | 0.0013 | **>0.9999** | **ns** |

| | | | |
|---|---|---|---|
| **SN** | | | |
| Saline vs. Ethanol-1.2g/kg | -0.0851 | **0.9581** | **ns** |
| Saline vs. GiGA1-30mg/Kg | -0.2705 | **0.3915** | **ns** |
| Saline vs. GiGA1+Ethanol | -0.5175 | **0.0186** | ● |

| | | | |
|---|---|---|---|
| **APT** | | | |
| Saline vs. Ethanol-1.2g/kg | -0.2447 | **0.4798** | **ns** |
| Saline vs. GiGA1-30mg/Kg | -0.4189 | **0.0783** | **ns** |
| Saline vs. GiGA1+Ethanol | -0.4759 | **0.0353** | ● |

| | | | |
|---|---|---|---|
| **PrEW** | | | |
| Saline vs. Ethanol-1.2g/kg | 1.0150 | **<0.0001** | ●●●● |
| Saline vs. GiGA1-30mg/Kg | -0.4144 | **0.0830** | **ns** |
| Saline vs. GiGA1+Ethanol | -0.0345 | **0.9970** | **ns** |

**Table S8 (relates to Figure S1A)**

GiGA1 treatment in Cocaine-CPP (Time spent in Paired side)

| **Females** | **N=10** | | ***Post-hoc Comparison*** | ***Holm-Šídák's*** |
|---|---|---|---|---|
| ***2way-ANOVA*** | *F (DFn, DFd)* | *P value* | **Pre-test vs Post-test** | *Adjusted P Value* |

| | | | | | | |
|---|---|---|---|---|---|---|
| Treatment x Test | F (1, 18) = 0.1628 | **0.6914** | **ns** | | | |
| Treatment | F (1, 18) = 0.03547 | **0.8527** | **ns** | Vehicle + Cocaine | **0.0370** | * |
| Test | F (1, 18) = 12.88 | **0.0021** | **##** | GiGA1 + Ethanol | **0.0224** | * |

**Table S9 (relates to Figure S1B)**

GiGA1 treatment in NOR (Discrimination index)

| **Females** | **N=10** | | | ***Post-hoc Comparison*** | ***Holm-Šídák's*** | |
|---|---|---|---|---|---|---|
| ***2way-ANOVA*** | *F (DFn, DFd)* | *P value* | | **Pre-test vs Post-test** | *Adjusted P Value* | |
| Treatment x Test | F (1, 14) = 0.8177 | **0.3811** | **ns** | Vehicle-group | **0.0048** | ** |
| Test | F (1, 14) = 18.67 | **0.0007** | **###** | GiGA1-group | **0.0299** | * |

**Table S10 (relates to Figure S2)**

Dose-response

| | **N=12** | | | ***Post-hoc Comparison*** | ***Šídák's*** | |
|---|---|---|---|---|---|---|
| ***1way-ANOVA*** | *F (DFn, DFd)* | *P value* | | **Vehicle vs Dose** | *Adjusted P Value* | |
| | | | | GiGA1-20mg/kg | 0.9841 | **ns** |
| | F (3, 44) = 16.88 | P<0.0001 | **####** | GiGA1-30mg/kg | 0.0001 | *** |
| | | | | GiGA1-40mg/kg | <0.0001 | **** |

**Table S11 (relates to Figure S3)**

Intake Baclofen treatment (Males)

| ***Paired t-test*** | *Group* | *N* | *t, df* | *P value* | |
|---|---|---|---|---|---|
| Baclofen-7.5mg/kg vs Vehicle | Ethanol | 16 | t=6.462, df=15 | **<0.0001** | **** |

| ***Paired t-test*** | *Group* | *N* | *t, df* | *P value* | |
|---|---|---|---|---|---|
| Baclofen-7.5mg/kg vs Vehicle | Water | 15 | t=1.517, df=14 | **0.1514** | **ns** |

Preference Baclofen treatment (Males)

| ***Paired t-test*** | *Group* | *N* | *t, df* | *P value* | |
|---|---|---|---|---|---|
| Baclofen-7.5mg/kg vs Vehicle | Ethanol | 16 | t=1.128, df=15 | **0.2770** | **ns** |

BAC Baclofen treatment

| ***Paired t-test*** | *Group* | *N* | *t, df* | *P value* | |
|---|---|---|---|---|---|
| Baclofen-7.5mg/kg vs Vehicle | males | 16 | t=4.337, df=15 | **0.0006** | *** |

Sucrose intake Baclofen treatment

| ***Paired t-test*** | *Group* | *N* | *t, df* | *P value* | |
|---|---|---|---|---|---|
| Baclofen-7.5mg/kg vs Vehicle | males | 16 | t=3.528, df=15 | **0.0030** | ** |
